# An Engineered Antibody with Broad Protective Efficacy in Murine Models of SARS and COVID-19

**DOI:** 10.1101/2020.11.17.385500

**Authors:** C. Garrett Rappazzo, Longping V. Tse, Chengzi I. Kaku, Daniel Wrapp, Mrunal Sakharkar, Deli Huang, Laura M. Deveau, Thomas J. Yockachonis, Andrew S. Herbert, Michael B. Battles, Cecilia M. O’Brien, Michael E. Brown, James C. Geoghegan, Jonathan Belk, Linghang Peng, Linlin Yang, Trevor D. Scobey, Dennis R. Burton, David Nemazee, John M. Dye, James E. Voss, Bronwyn M. Gunn, Jason S. McLellan, Ralph S. Baric, Lisa E. Gralinski, Laura M. Walker

**Affiliations:** Adimab LLC, Lebanon, NH 03766, USA; Department of Epidemiology, The University of North Carolina at Chapel Hill, Chapel Hill, NC 27599, USA; Department of Molecular Biosciences, The University of Texas at Austin, Austin, TX 78712, USA; Department of Immunology and Microbiology, The Scripps Research Institute, La Jolla, CA 92037, USA; Paul G. Allen School of Global Animal Health, Washington State University, Pullman, WA 99164, USA; U.S. Army Medical Research Institute of Infectious Diseases, Frederick, MD 21702, USA; The Geneva Foundation, 917 Pacific Avenue, Tacoma, WA 98402, USA; IAVI Neutralizing Antibody Center, The Scripps Research Institute, La Jolla, CA 92037, USA; Consortium for HIV/AIDS Vaccine Development (CHAVD), The Scripps Research Institute, La Jolla, CA 92037, USA; Ragon Institute of Massachusetts General Hospital, Massachusetts Institute of Technology, and Harvard, Cambridge, MA 02139, USA; Departments of Microbiology and Immunology, The University of North Carolina at Chapel Hill, Chapel Hill, NC 27599, USA; Adagio Therapeutics, Inc., Waltham, MA 02451, USA

**Author notes:** Corresponding authors. (R.S.B.); (L.E.G.); (L.M.W.). These authors contributed equally to this work (C. Garrett Rappazzo, Longping V. Tse).

## Abstract

The recurrent zoonotic spillover of coronaviruses (CoVs) into the human population underscores the need for broadly active countermeasures. Here, we employed a directed evolution approach to engineer three SARS-CoV-2 antibodies for enhanced neutralization breadth and potency. One of the affinity-matured variants, ADG-2, displays strong binding activity to a large panel of sarbecovirus receptor binding domains (RBDs) and neutralizes representative epidemic sarbecoviruses with remarkable potency. Structural and biochemical studies demonstrate that ADG-2 employs a unique angle of approach to recognize a highly conserved epitope overlapping the receptor binding site. In murine models of SARS-CoV and SARS-CoV-2 infection, passive transfer of ADG-2 provided complete protection against respiratory burden, viral replication in the lungs, and lung pathology. Altogether, ADG-2 represents a promising broad-spectrum therapeutic candidate for the treatment and prevention of SARS-CoV-2 and future emerging SARS-like CoVs.

---

Over the past two decades, three pathogenic CoVs have emerged from zoonotic reservoirs to cause outbreaks of deadly pneumonia in humans: severe acute respiratory syndrome coronavirus (SARS-CoV), Middle-East respiratory syndrome coronavirus (MERS-CoV), and severe acute respiratory syndrome coronavirus-2 (SARS-CoV-2) (*1*). SARS-CoV emerged in 2002 in the Guangdong province of China and infected ~8000 people with a case fatality rate of ~10% before being contained by public health measures (*2*). MERS-CoV emerged in the human population in 2012 and is still a significant public health threat in the Middle East (*3, 4*). In late 2019, SARS-CoV-2 emerged in the city of Wuhan in China’s Hubei province and rapidly caused an ongoing pandemic that has resulted in over a million deaths while disrupting the global economy (*1, 5*). Currently, there are no approved vaccines to prevent SARS-CoV-2 infection and only one antiviral drug has been approved to treat SARS-CoV-2 associated disease (i.e. COVID-19). Furthermore, the recurrent zoonotic spillover of CoVs into the human population, along with the broad diversity of SARS-like CoVs circulating in animal reservoirs (*6*), suggests that novel pathogenic CoVs are likely to emerge in the future and underscores the need for broadly active countermeasures.

Similar to other CoVs, the SARS-CoV-2 spike (S) protein mediates viral entry and is the only known target for neutralizing antibodies (nAbs). The S glycoprotein consists of two functional subunits, S1 and S2, that mediate receptor binding and viral fusion, respectively. Previous studies have shown that the vast majority of potent neutralizing antibodies induced by natural CoV infection target the RBD on the S1 subunit (*7–10*). Although SARS-CoV and SARS-CoV-2 both belong to the sarbecovirus subgenus and their S glycoproteins share 76% amino acid identity, only a handful of cross-neutralizing antibodies have been described to date (*11–13*). These rare broadly neutralizing antibodies (bnAbs) represent an attractive opportunity for therapeutic drug stockpiling to prevent or mitigate future outbreaks of SARS-related CoVs, but their limited neutralization potency may translate into suboptimal protective efficacy or impractical dosing regimens. Here, we show that such bnAbs can be engineered for improved neutralization potency while retaining neutralization breadth, and we demonstrate that these bnAbs can provide broad protection *in vivo*.

We recently isolated several antibodies from the memory B cells of a 2003 SARS survivor that cross-neutralize multiple SARS-related viruses with relatively modest potency (*11*). Although breadth and potency are often opposing characteristics, we sought to engineer these bnAbs for improved neutralization potency against SARS-CoV-2, while also maintaining or improving neutralization breadth and potency against other SARS-related viruses. Because binding affinity and neutralization potency are generally well-correlated (*14*), we employed yeast-surface display technology to improve the binding affinities of three of the bnAbs (ADI-55688, ADI-55689, and ADI-56046) for a prefusion-stabilized SARS-CoV-2 S protein (*11, 15–17*).

Yeast display libraries were generated by introducing diversity into the heavy (HC)- and light-chain (LC) variable genes of ADI-55688, ADI-55689, and ADI-56046 through oligonucleotide-based mutagenesis and transformation into *Saccharomyces cerevisiae* by homologous recombination (*15*). Following four rounds of selection with a recombinant SARS-CoV-2 S1 protein, improved binding populations were sorted, and between 20 and 50 unique clones from each lineage were screened for binding to SARS-CoV-2 S (*17*) (Fig. 1A, B and Fig. S1). The highest affinity binders from each of the three lineages bound to the SARS-CoV-2 S protein with monovalent equilibrium dissociation constants (K_D_s) in the picomolar range, representing 25 to 630-fold improvements in binding relative to their respective parental clones (Fig. 1B and Fig. S2). To determine whether the improvements in SARS-CoV-2 S binding affinity translated into enhanced neutralization potency, we selected between 9 and 14 affinity-matured progeny from each lineage, and evaluated them for SARS-CoV-2 neutralizing activity in a murine leukemia virus (MLV) pseudovirus assay (*18*). We also measured the neutralizing activities of several clinical-stage neutralizing antibodies (nAbs) (S309, REGN10933, REGN10987, and CB6/LY-CoV016) as benchmarks (*12, 19, 20*). All of the affinity-matured antibodies showed improved neutralizing activity relative to their parental clones, and the most potent neutralizers from each lineage (ADG-1, ADG-2, and ADG-3) displayed neutralization IC_50_s that were comparable to or lower than those observed for the clinical SARS-CoV-2 nAb controls (Fig. 1B).

**Figure 1.**
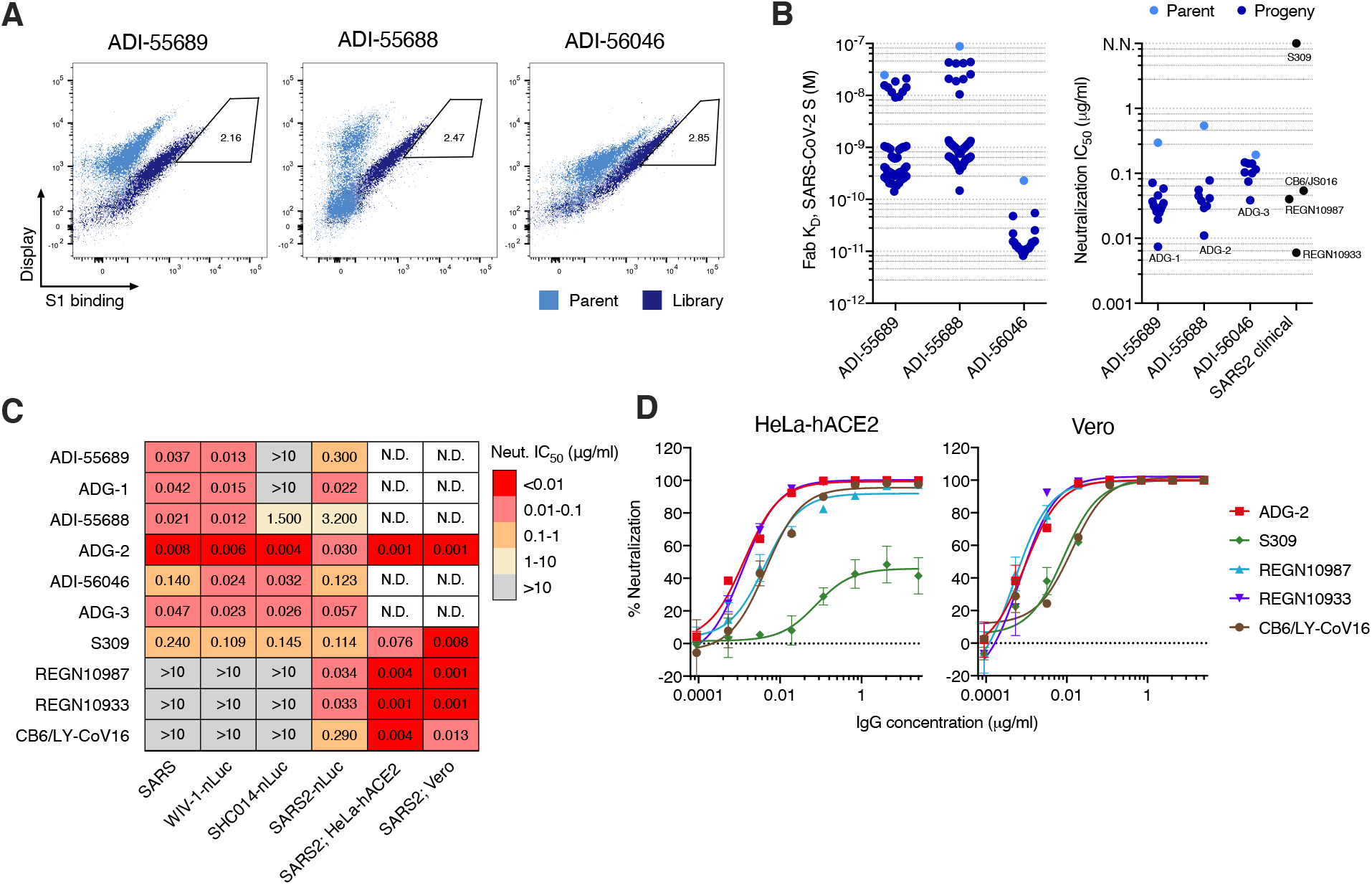
Engineering of SARS-CoV-2 antibodies for enhanced neutralization breadth and potency. **(A)** Flow cytometry plots from the terminal round of selection showing binding of parental antibodies (light blue) and affinity maturation library antibodies (dark blue) to the SARS-CoV-2 S1 protein at 1 nM. Gates indicate the yeast populations sorted for antibody sequencing and characterization. **(B)** Dot plots of Fab binding affinities (left) and MLV-SARS-CoV-2 pseudovirus neutralization IC_50_s (right) of parental antibodies and affinity matured progeny. SARS-CoV-2 clinical antibodies are shown for comparison. **(C)** Heat map showing the neutralization IC_50_s of the indicated antibodies against authentic SARS-CoV, WIV-1-nLuc, SHC014-nLuc, SARS-CoV-2-nLuc, and SARS-CoV-2 using either HeLa-hACE2 or Vero target cells. SARS-CoV assays were performed on Vero cells. WIV-1-nLuc, SHCO14-nLuc, and SARS-CoV-2 nLuc assays were performed on Vero cells with recombinant, reverse genetics-derived viruses encoding a nano-luciferase reporter gene. **(D)** Authentic SARS-CoV-2 neutralization titrations performed using either HeLa-hACE2 (left) or Vero (right) target cells. The curves were fit by nonlinear regression. Error bars represent standard deviation. N.D., not determined; N.N., non-neutralizing.

Because *in vitro* engineering can lead to polyspecificity with potential risks of off-target binding and accelerated clearance *in vivo (21)*, we assessed the polyspecificity of ADG-1, ADG-2, and ADG-3 using a previously described assay that has been shown to be predictive of serum half-life in humans (*22*). All three antibodies lacked polyreactivity in this assay, indicating a low risk for poor pharmacokinetic behavior (Fig. S3). The three antibodies also showed low hydrophobicity, a low propensity for self-interaction, and thermal stabilities within the range observed for clinically approved antibodies (Fig. S3). In summary, the process of *in vitro* engineering did not negatively impact biophysical properties that are often linked to down-stream behaviors such as serum half-life, ease of manufacturing, ability to formulate to high concentrations, and long-term stability.

To determine whether the process of SARS-CoV-2 affinity engineering impacted neutralization breadth, we evaluated ADG-1, ADG-2, and ADG-3, as well as their respective parental antibodies, for neutralizing activity against a panel of representative authentic clade I sarbecoviruses (SARS-CoV, SHC014, SARS-CoV-2, and WIV-1). Consistent with the MLV-SARS-CoV-2 assay results, ADG-2 displayed highly potent neutralizing activity against authentic SARS-CoV-2, with an IC_50_ comparable to or lower than that observed for the benchmark SARS-CoV-2 nAbs (Fig. 1C and Fig. S4). Furthermore, in contrast to the benchmark nAbs, ADG-2 displayed high neutralization potency against SARS-CoV and the two SARS-related bat viruses, with IC_50_s between 4 and 8 ng/mL (Fig. 1C and Fig. S4). ADG-3 and the clinical nAb S309 also cross-neutralized all four sarbecoviruses, but with markedly lower potency than ADG-2. Finally, ADG-1 potently neutralized SARS-CoV-2, SARS-CoV, and WIV1, but it lacked activity against SHC014.

Based on its potent cross-neutralization and favorable biophysical properties, we selected ADG-2 as a lead therapeutic candidate and confirmed its potent neutralizing activity in two alternative authentic SARS-CoV-2 neutralization assays (IC_50_ ~1 ng/mL) (Fig. 1C, D and Fig. S4). Interestingly, ADG-2, CB6/LY-CoV016, REGN10987 and REGN10933 reached 100% neutralization on both Vero and HeLa-hACE2 target cells in this assay, whereas S309 showed complete neutralization on Vero target cells but plateaued at approximately 40% neutralization on HeLa-hACE2 target cells (Fig. 1D). S309 also failed to neutralize MLV-SARS-CoV-2 on HeLa-hACE2 target cells (Fig. 1B). The reason for this is unclear but may relate to glycan heterogeneity within the S309 epitope (*12*) coupled with differences in receptor expression or protease cleavage efficiency between the two types of target cells (*23*). Because SARS-CoV-2 D614G has emerged as the dominant pandemic strain (*24*), we also evaluated ADG-2 for neutralizing activity against this variant in the MLV pseudovirus assay. As expected, based on the location of the D614G substitution outside of the RBD, ADG-2 neutralized the D614G variant with equivalent potency as wild-type (WT) SARS-CoV-2 (Fig. S5).

We further assessed the breadth of sarbecovirus recognition by ADG-2 by measuring its apparent binding affinity (K_D_^App^) to a panel of 17 representative sarbecovirus RBDs expressed on the surface of yeast (*25*). Thirteen viruses were selected from clade I — representing the closest known relatives of SARS-CoV-2 (GD-Pangolin and RaTG13) to the most divergent (SHC014 and Rs4231) — as well as four viruses from the distantly related clades 2 and 3, which do not utilize ACE2 as a host receptor (*26*) (Fig. 2A). Recombinant hACE2-Fc and the benchmark SARS-CoV-2 nAbs described above were also included as controls. Consistent with previous reports (*17, 25*), hACE2 only recognized clade I RBDs and bound with higher affinity to SARS-CoV-2 than SARS-CoV (Fig. 2B). In addition, the benchmark SARS-CoV-2 nAbs CB6/LY-CoV016, REGN10987, and REGN10933 bound to the SARS-CoV-2 RBD with K_D_^Apps^ comparable to published reports (Fig. 2B) (*19, 20*). Notably, S309 displayed diminished binding in this expression platform, likely due to recognition of an epitope containing an N-glycan that may be hyper-mannosylated in yeast (*12*).

**Figure 2.**
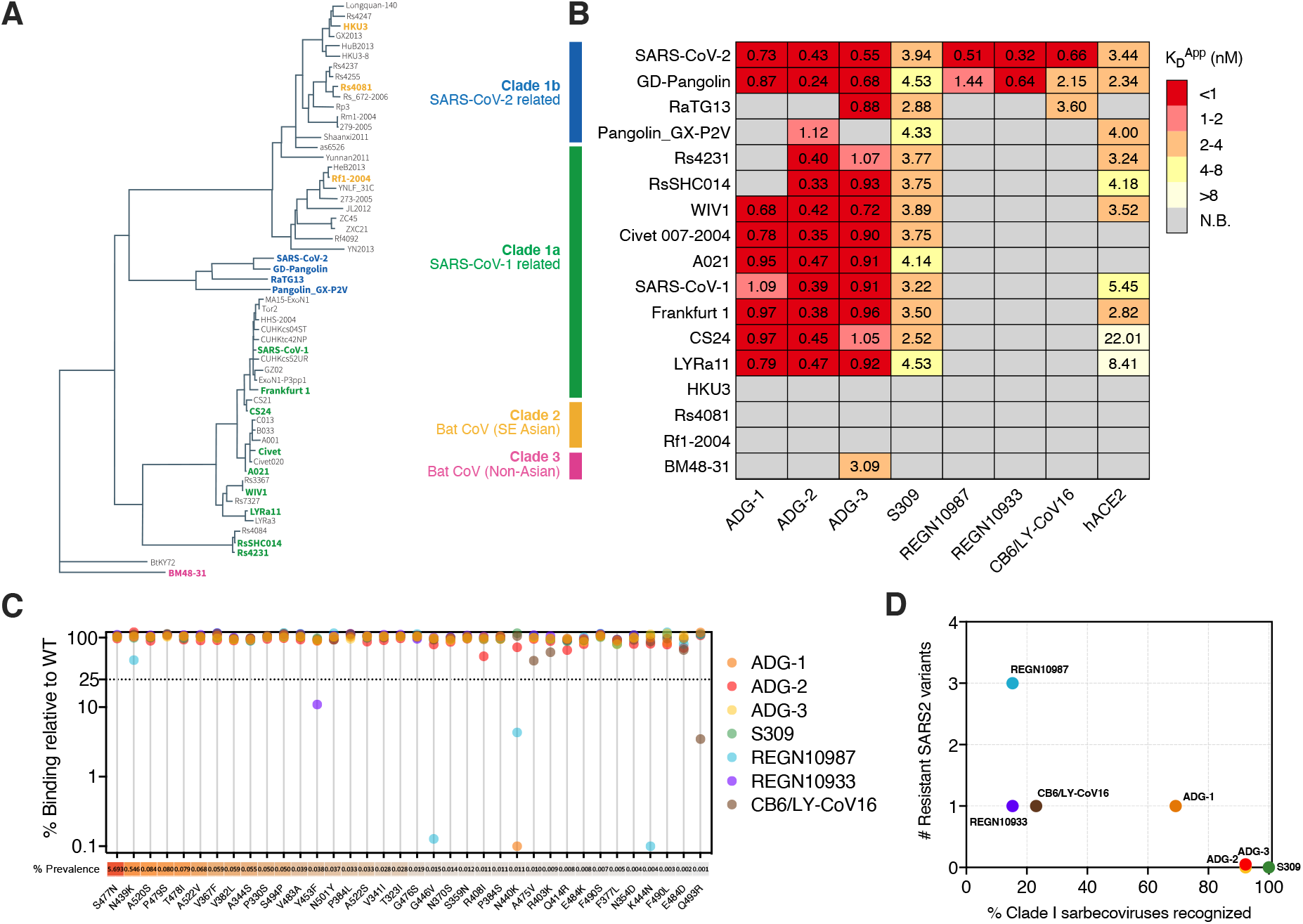
Breadth of antibody binding to diverse sarbecoviruses and circulating SARS-CoV-2 variants. **(A)** Phylogenetic tree of 57 sarbecoviruses constructed via MAFFT and maximum likelihood analysis of RBD-SD1 amino acid sequences extracted from the European Nucleotide Archive and GISAID database. Representative sarbecovirus RBDs selected for further study are denoted in bold and colored according to their canonical phylogenetic lineages. **(B)** Heat map of antibody and recombinant hACE2 binding to yeast-displayed RBDs from 17 representative sarbecoviruses, grouped by phylogenetic lineages. K_D_^App^ values were calculated by normalized nonlinear regression fitting. **(C)** Antibody binding to naturally-occurring SARS-CoV-2 RBD variants displayed on the surface of yeast. SARS-CoV-2 sequences were retrieved from the GISAID database on July 14, 2020 (*n* = 63551). Antibody binding signal was normalized to RBD expression and calculated as percent binding of the variant relative to the WT SARS-CoV-2 RBD, assessed at their respective K_D_^App^ concentrations for the WT construct. The prevalence of each variant, calculated from deposited sequences on October 19, 2020 (*n* = 148115), is shown as a percentage of the total number of sequences analyzed. **(D)** Correlation between the number of resistant SARS-CoV-2 variants and percentage of clade I sarbecovirus RBDs recognized. N.B., non-binder.

Consistent with their broadly neutralizing activities, S309, ADG-2, and ADG-3 displayed remarkably broad binding reactivity to clade I sarbecovirus RBDs, with ADG-2 and ADG-3 strongly binding 12/13 viruses and S309 binding all 13 (Fig. 2B). In contrast, ADG-1 only bound to 9/13 viruses and CB6/LY-CoV016, REGN10987, and REGN10933 bound only the closest evolutionary neighbor(s) of SARS-CoV-2, consistent with their narrow neutralization profiles (Fig. 2B and Fig. 1C). Importantly, ADG-2 bound with high affinity (K_D_^App^ 0.24-1.12 nM) to every clade I sarbecovirus RBD that exhibited detectable hACE2 binding in our assay. This finding supports the high degree of ADG-2 epitope conservation among sarbecoviruses that use hACE2 as a receptor.

Several recent studies have shown that RBD mutants that are resistant to commonly elicited SARS-CoV-2 nAbs are circulating at low levels in the human population (*24, 27*). We therefore sought to assess the breadth of ADG-2 binding to naturally circulating SARS-CoV-2 variants that contain single point mutations in the RBD. ADG-1, ADG-3, and the benchmark SARS-CoV-2 nAbs were also included as comparators. Using the yeast surface-display platform described above, we expressed the 30 most frequently observed SARS-CoV-2 RBD variants reported in the GISAID database as well as six naturally circulating SARS-CoV-2 variants that have been shown to be resistant to previously described SARS-CoV-2 nAbs (*24, 27, 28*). One or more of the 36 SARS-CoV-2 variants exhibited loss of binding to ADG-1, CB6/LY-CoV016, REGN10987, and REGN10933, as defined by >75% loss relative to the WT construct (Fig. 2C). Notably, the loss-of-binding variants identified for REGN10987 and REGN10933 partially overlapped with those identified in previous *in vitro* neutralization escape studies, validating the use of RBD display for the prediction of antibody escape mutations (*29*). In contrast, ADG-2, ADG-3, and S309 bound to all 36 variants at levels ≥50% of WT SARS-CoV-2 (Fig. 2C). This result, combined with the remarkable neutralization breadth observed for these three mAbs (Fig. 1C and Fig. 2B, D), suggests a potential link between epitope conservation and resistance to viral escape.

To gain further insight into the antigenic surface recognized by ADG-2, we generated a mutagenized yeast surface-display RBD library and performed rounds of selection to identify RBD variants that exhibited loss of binding to ADG-2 relative to the WT construct (Fig. 3A, Fig. S6A, B). To exclude mutations that globally disrupt the conformation of the RBD, a final round of positive selection was performed using a mixture of recombinant hACE2 and two RBD-directed mAbs (S309 and CR3022) that target non-overlapping epitopes distinct from the ADG-2 binding site (*12, 30*) (Fig. S6B, and Fig. S7). Selected RBD mutants encoding single amino acid substitutions were individually tested for binding to ADG-2, recombinant hACE2, CR3022, and S309 to confirm site-specific knock-down mutations (Fig. S6C). Substitutions at only four RBD positions specifically abrogated ADG-2 binding: D405E, G502E/R/V, G504A/D/R/S/V and Y505C/N/S (Fig. 3B). These four residues are highly conserved among the clade I sarbecovirus subgenus and invariant among SARS-CoV-1, SARS-CoV-2, SHC014 and WIV1 viruses (Fig. 3C), providing a molecular explanation for the breadth of binding and neutralization exhibited by ADG-2. Consistent with the conservation of these residues among clade I sarbecoviruses, none of the substitutions that impacted ADG-2 binding were present in full-length SARS-CoV-2 sequences deposited in the GISAID database as of October 19, 2020. In addition, 3 of the 4 identified mutations that abrogate ADG-2 binding lie within the hACE2 binding site (*31*) and at least one mutation at each position (G502E/R/V, G504V and Y505C/N/S) also abrogated hACE2 binding (Fig. 3B), likely accounting for their absence among circulating SARS-CoV-2 isolates. These results suggest that the evolutionary conservation of the ADG-2 epitope is likely directly linked to ACE2 binding.

**Figure 3.**
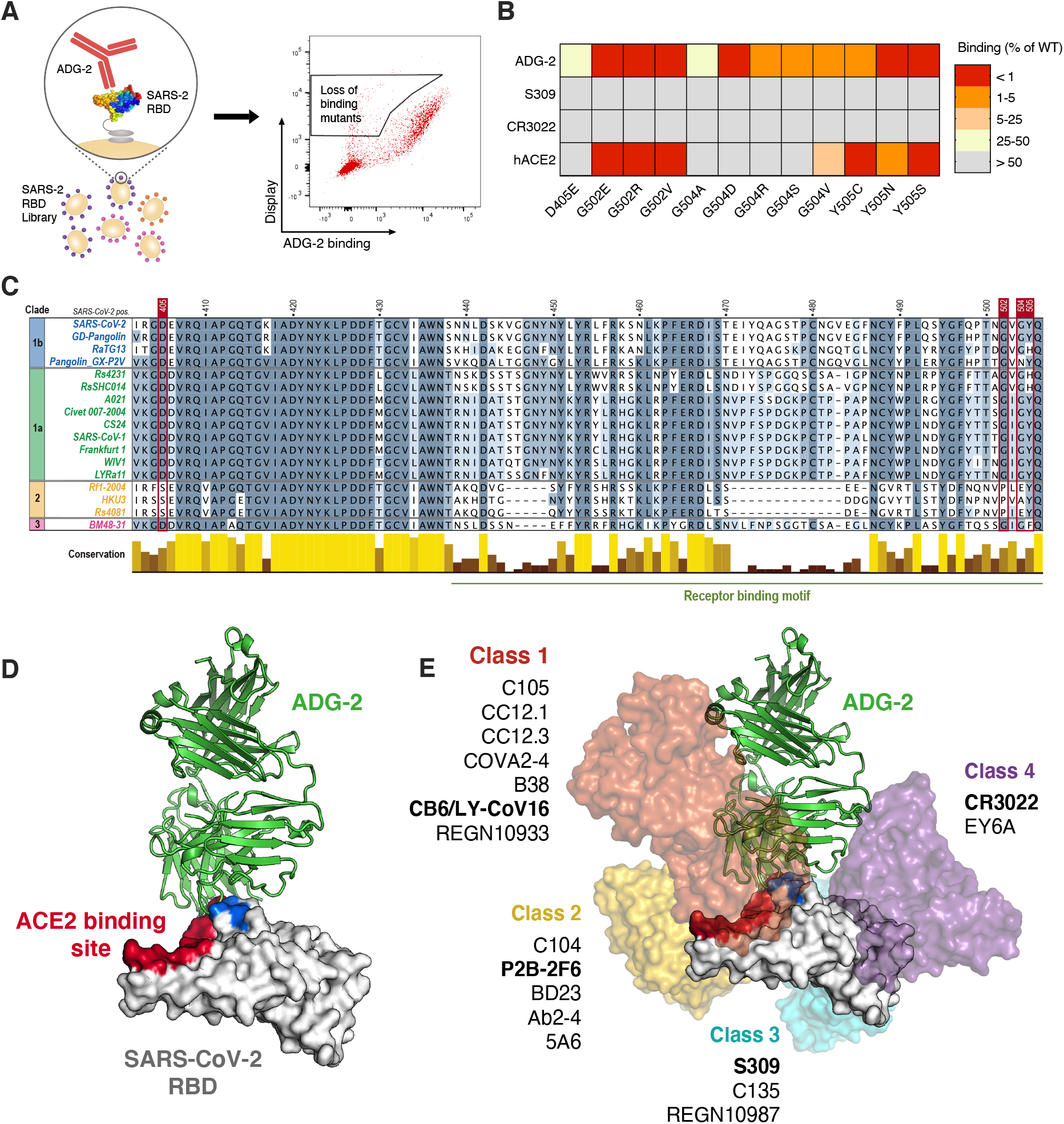
ADG-2 binds to an evolutionarily conserved epitope on the SARS-CoV-2 RBD overlapping with the hACE2 binding site. **(A)** Schematic showing the generation and selection of a mutagenized, yeast surface-displayed SARS-CoV-2 RBD library to identify mutations that knock-down ADG-2 binding. **(B)** Heat map showing mutations that abrogate binding of ADG-2 to the SARS-CoV-2 RBD. S309 and CR3022, which bind non-overlapping epitopes distinct from the ADG-2 binding site, are included to control for mutations that globally disrupt the conformation of the RBD. Values indicate percent antibody or recombinant hACE2-Fc binding to the mutant SARS-CoV-2 RBD relative to the WT SARS-CoV-2 RBD, assessed at their respective EC_80_ concentrations for the WT RBD construct. **(C)** Protein sequence alignment of representative sarbecovirus RBDs with sequences colored by percentage sequence identity and conservation shown as a bar plot. Positions delineating the receptor binding motif are based on the SARS-CoV-2 RBD. Residues determined to be important for ADG-2 binding based on the data shown in (B) are denoted in red. **(D)** Cryo-EM reconstruction of the SARS-CoV-2 RBD bound by ADG-2, with ADG-2 knock-down mutations and the hACE2 binding site highlighted in blue and red, respectively. **(E)** Structures of previously reported antibodies (bold) representing frequently observed SARS-CoV-2 nAb classes 1-4 overlaid on the ADG-2 structure (D), with additional representative SARS-CoV-2 nAbs listed.

To support the results of this experiment, we performed low-resolution cryogenic electron microscopy (cryo-EM) of the complex of ADG-2 bound to prefusion-stabilized SARS-CoV-2 S. This yielded a ~6Å resolution 3D reconstruction that clearly had at least one ADG-2 Fab bound to an RBD in the up conformation and allowed us to unambiguously dock in previously determined high-resolution models of the SARS-CoV-2 spike and a homologous Fab (Fig. 3D, Fig. S8A-D, Table S1). Consistent with our fine epitope mapping and competitive binding experiments (Fig. 3B and Fig. S7C), the epitope recognized by ADG-2 overlaps with the hACE2-binding site and each position identified by epitope mapping clustered to the cleft between the heavy and light chains of ADG-2 (Fig. 3D). This epitope also partially overlaps with those recognized by frequently observed “class 1” SARS-CoV-2 nAbs, which are exemplified by *VH3-53* antibodies with short CDRH3s and compete with hACE2 in the RBD “up” conformation (*32*) (Fig. 3E). However, in contrast to previously reported nAbs in this class, ADG-2 binds with a divergent angle of approach and displays broadly neutralizing activity (*32*) (Fig. 3E, Fig. 1C, and Fig. S8E). Thus, ADG-2 binds to a highly conserved motif via a unique angle of approach, providing additional structural insight into its broad recognition of SARS-like CoVs.

Because Fc-mediated effector functions can contribute to protection independently of viral neutralization, we next assessed the ability of ADG-2 to induce antibody-dependent natural killer cell activation and degranulation (ADNKDA), antibody-dependent cellular phagocytosis (ADCP) mediated by monocytes and neutrophils, and antibody-mediated complement deposition (ADCD) using previously described *in vitro* assays (*33*). Benchmark SARS-CoV-2 nAbs S309 and REGN10987 were also included as comparators. ADG-2 displayed a highly polyfunctional profile, resulting in the induction of phagocytosis by monocytes and neutrophils, deposition of the complement component C3, and induction of NK cell degranulation (a surrogate marker of ADCC) and activation (Fig. 4). Interestingly, while ADG-2, S309, and REGN10957 showed comparable recruitment of phagocytosis (Fig. 4B), these antibodies differed with respect to complement deposition and NK cell activation (Fig. 4A, C); S309 showed reduced complement deposition compared with ADG-2 and REGN10987, and ADG-2 showed superior NK cell activation over both S309 and REGN10987 (Fig.4). In summary, ADG-2 robustly triggers diverse Fc-mediated effector activities with potencies comparable or superior to those of current lead SARS-CoV-2 clinical antibodies. However, it should be noted that the contribution of extra-neutralizing activities to protection against SARS-CoV-2 is currently unknown and may vary among different antibody specificities.

**Figure 4.**
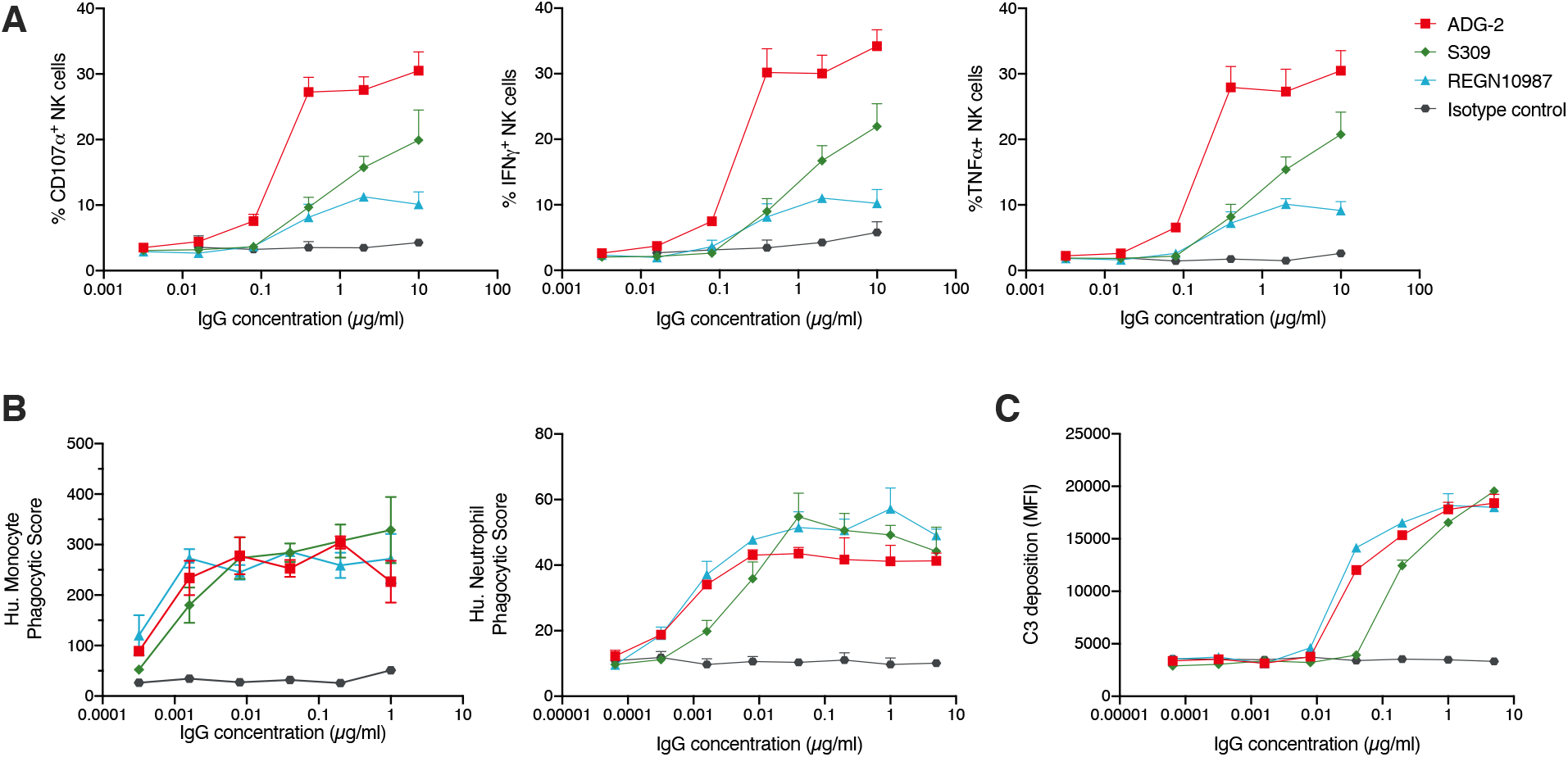
ADG-2 triggers Fc-mediated effector functions. The indicated antibodies were assessed for the ability to induce Fc-mediated effector functions against RBD-coated targets at varying concentrations. **(A)** Primary human NK cells were analyzed for surface expression of CD107a, indicating degranulation (left), and the production of IFNγ (middle) or TNFα (right) following incubation with antibody-RBD immune complexes for 5 hours. **(B)** Antibody-mediated phagocytosis of RBD-coated fluorescent beads by differentiated HL-60 neutrophils (left) or THP-1 monocytes (right) was measured following incubation with immune complexes for 18 hours. **(C)** Antibody-mediated complement deposition was measured by detection of complement component C3 onto RBD-coated fluorescent beads following incubation of guinea pig complement with immune complexes for 20 minutes.

Finally, we tested the ability of ADG-2 to provide broad *in vivo* protection in immunocompetent mouse models of SARS and COVID-19 using mouse-adapted SARS-CoV (MA15)- and SARS-CoV-2 (MA10), respectively (*34, 35*). Balb/c mice were prophylactically treated with either 200 *>μ*g of ADG-2 or PBS via IP injection 12 hours prior to intranasal challenge with a 10^3^ PFU dose of MA15 or MA10. All mice were monitored daily for weight loss and changes in respiratory function and groups of mice were euthanized at day two or four post-infection to allow for measurement of virus replication in the lung and analysis of lung histopathology. We observed substantial, progressive weight loss in sham-treated mice infected with both viruses along with increases in Penh, a calculated measure of airway resistance (*35*). In contrast, mice treated prophylactically with ADG-2 demonstrated minimal weight loss, no change in Penh and no signs of gross pathology at the time of harvest (Fig. 5A, B). Furthermore, prophylactic antibody treatment prevented viral replication in the lungs at both two and four days post-infection (dpi). We next investigated the ability of ADG-2 to act anti-virally against SARS-CoV-2 MA10 in a therapeutic setting. Mice were treated with 200 *>μ*g of ADG-2 or PBS 12 hours following intranasal challenge with a 10^3^ PFU dose of MA10. Mice given therapeutic ADG-2 had intermediate levels of weight loss, moderate respiratory function changes and some gross lung pathology; significantly more than prophylactically-treated mice but significantly less than sham-treated mice (Fig. 5C). Therapeutic antibody treatment also resulted in a significant reduction in lung viral loads at four dpi, but not at two dpi, relative to sham-treated mice. We conclude that ADG-2 treatment can reduce disease burden in mice infected with both SARS-CoV MA15 and SARS-CoV-2 MA10.

**Figure 5.**
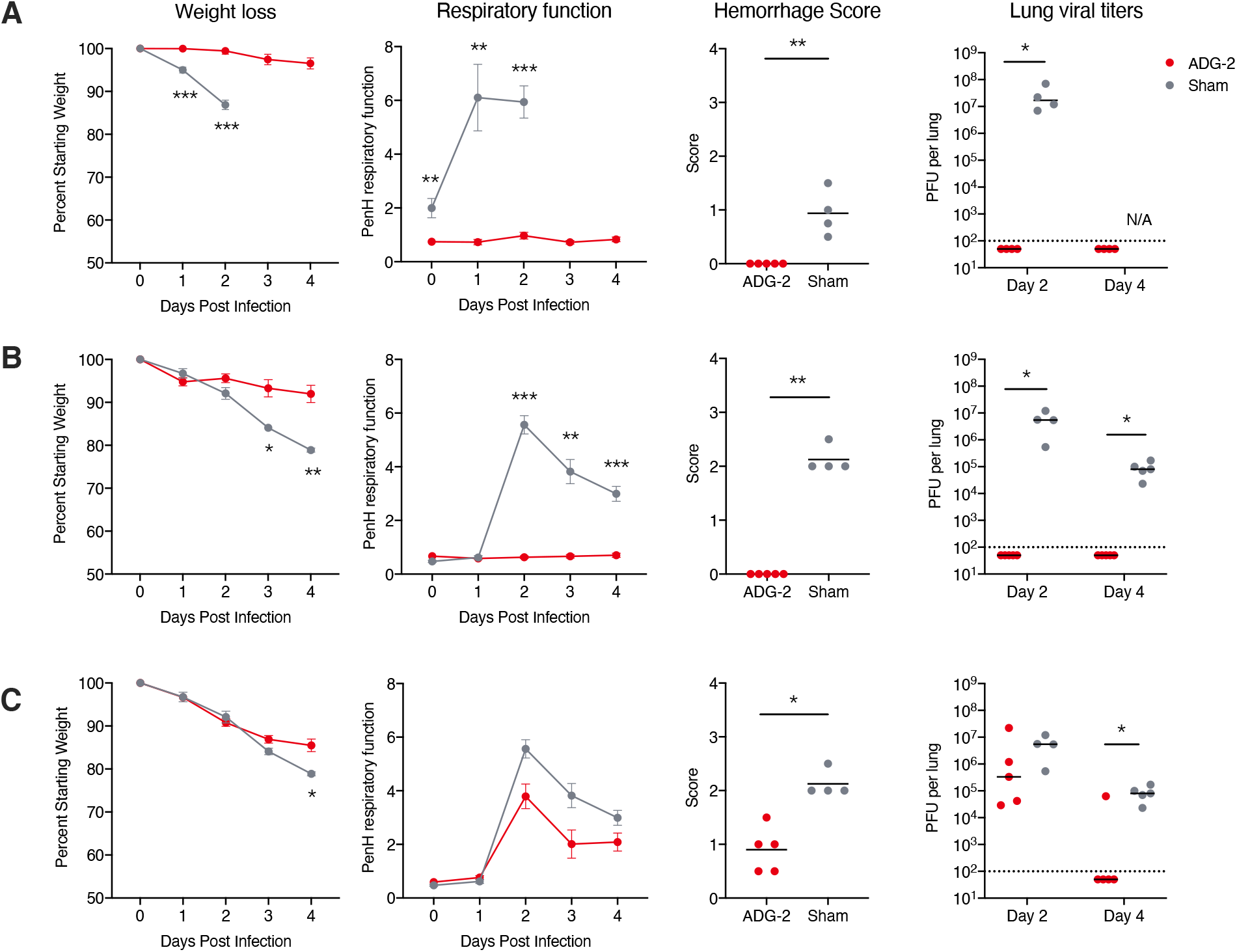
Prophylactic and therapeutic administration of ADG-2 protects mice from SARS-CoV- and SARS-CoV-2-associated viral diseases. Efficacy of prophylactic treatment with ADG-2 in **(A)** SARS-CoV-MA15 and **(B)** SARS-CoV-2-MA10 challenge models. A single dose of ADG-2 or sham treatment were delivered intraperitoneally 12 hours prior to infection. Mouse body weight and respiratory function were monitored for 4 days. Gross lung hemorrhage scores were determined on day 2 (MA15) or day 4 (MA10) post-infection and lung viral titers were measured on day 2 and day 4 post-infection. **(C)** Therapeutic treatment with ADG-2 or sham treatment at 12 hours post-SARS-CoV-2-MA10-infection. Mouse body weight, respiratory function, gross hemorrhage scores (day 2), and lung viral titers (days 2 and 4) were assessed as described above. Statistical comparisons were made using Mann-Whitney U tests or two-sided t-tests with Holm-Sidak corrections for multiple comparisons (*P < 0.05, **P < 0.01; ***P < 0.001). Dotted lines indicate the limit of detection.

Since the beginning of the COVID-19 pandemic, a plethora of potently neutralizing SARS-CoV-2 antibodies have been isolated, and some have rapidly advanced into clinical trials (*36*). However, the epitopes recognized by most of these nAbs are highly variable among other clade 1a and 1b sarbecoviruses, hence limiting their neutralization breadth and increasing their susceptibility to antibody escape mutations (*27*). Here, we described an engineered antibody that neutralizes SARS-CoV-2 with a potency that rivals current lead SARS-CoV-2 clinical nAbs, but also broadly neutralizes other clade I sarbecoviruses, potently triggers Fc-mediated effector functions, and provides significant protection against SARS and COVID-19 disease in mouse models. Thus, ADG-2 represents a promising candidate for the prevention and treatment of not only COVID-19 but also future respiratory diseases caused by pre-emergent SARS-related CoVs. Furthermore, our fine epitope mapping and structural studies demonstrate that ADG-2 employs a unique angle of approach to recognize a highly conserved epitope overlapping the receptor binding site. This epitope represents an Achilles’ heel for clade 1a and 1b sarbecoviruses and hence an attractive target for the rational design of “pan-SARS” vaccines that aim to elicit similar broadly protective antibodies.

## Acknowledgements

We thank T. Boland for assistance with SARS-CoV-2 sequence analysis and C. Williams for assistance with figure preparation. We thank E. Krauland and M. Vasquez for helpful comments on the manuscript. We thank J. Ludes-Meyers for assistance with cell transfection. All IgGs were sequenced by Adimab’s Molecular Core and produced by the High Throughput Expression group. BLI binding experiments were performed by Adimab’s Protein Analytics group. Opinions, conclusions, interpretations, and recommendations are those of the authors and are not necessarily endorsed by the U.S. Army. The mention of trade names or commercial products does not constitute endorsement or recommendation for use by the Department of the Army or the Department of Defense.

## Author contributions

L.M.W., L.E.G. and R.S.B conceived and designed the study. L.M.D. and J.B. performed the directed evolution experiments. L.V.T., D.H., A.S.H., C.M.O., L.P., L.Y., T.D.S., D.R.B., D.N., J.M.D., J.V. and R.S.B. developed, designed, and performed neutralization assays. M.E.B. and J.C.G. designed and supervised developability and biolayer interferometry assays. C.G.R., C.I.K., M.S., and M.B.B. designed and performed the yeast surface-display RBD experiments. D.W. and J.S.M. designed and performed Biacore SPR and structural assays. T.J.Y. and B.M.G. designed and performed Fc-effector functional assays. L.E.G. designed and performed the animal challenge studies. C.G.R., L.V.T., C.I.K., D.W., M.S., D.H., L.M.D., A.S.H., M.B.B., B.M.G., L.E.G., and L.M.W. analyzed the data. C.G.R., L.V.T., C.I.K., D.W., M.S. D.H., B.M.G., L.E.G., and L.M.W. wrote the manuscript and all authors reviewed and edited the paper.

## Data and material availability

IgGs are available from the corresponding author under MTA from Adagio Therapeutics, Inc.

## Supplementary Figures

**Figure S1.**
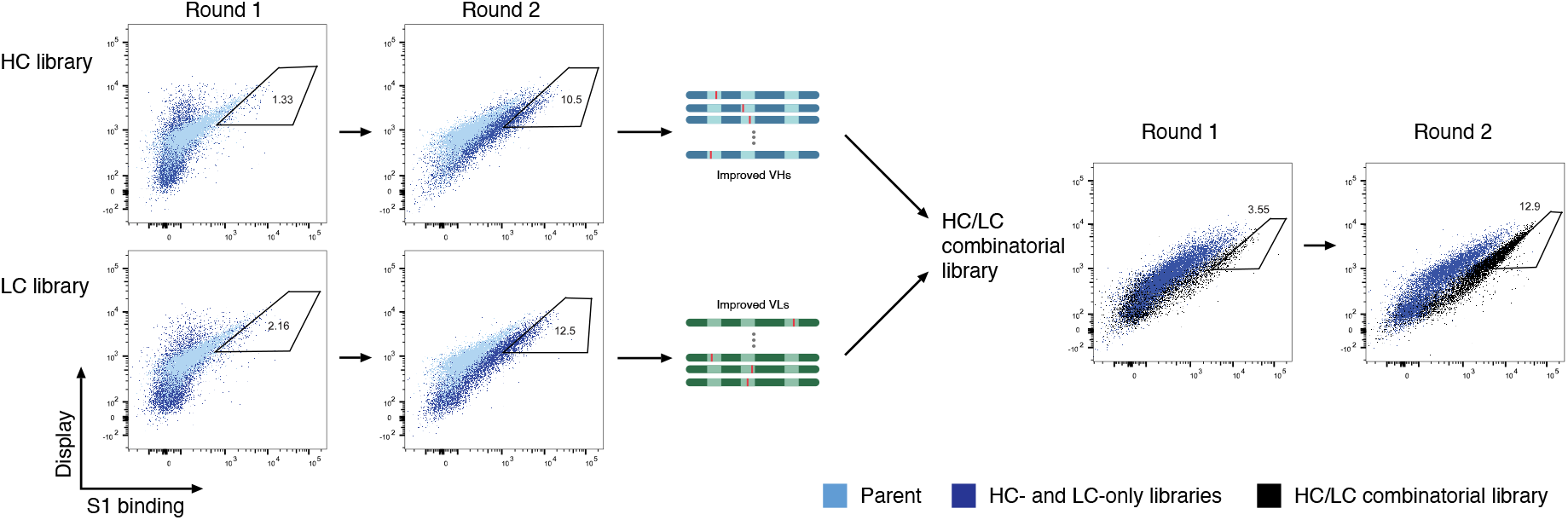
Representative selection strategy for affinity maturation libraries. **(A)** Flow cytometric sorting of libraries containing diversity in the HC (top) or LC (bottom) of ADI-55688. Libraries (dark blue) were sorted for improved binding to the SARS-CoV-2 S1 protein relative to the parent clone (light blue). Round 1 gates indicate the yeast populations that were sorted for a second round of selection, and round 2 gates indicate the yeast populations that were sorted for amplification of heavy- or light-chain variable region genes and subsequent transformation into yeast to generate a HC/LC combinatorial library. **(B)** Flow cytometric sorting of the HC/LC combinatorial library (black) for improved binding to the SARS-CoV-2 S1 protein relative to the round 2 output of the HC diversity libraries (dark blue). The round 1 gate indicates the yeast population that was sorted for a second round of selection and the round 2 gate indicates the yeast population that was sorted for individual colony isolation and sequencing.

**Figure S2.**
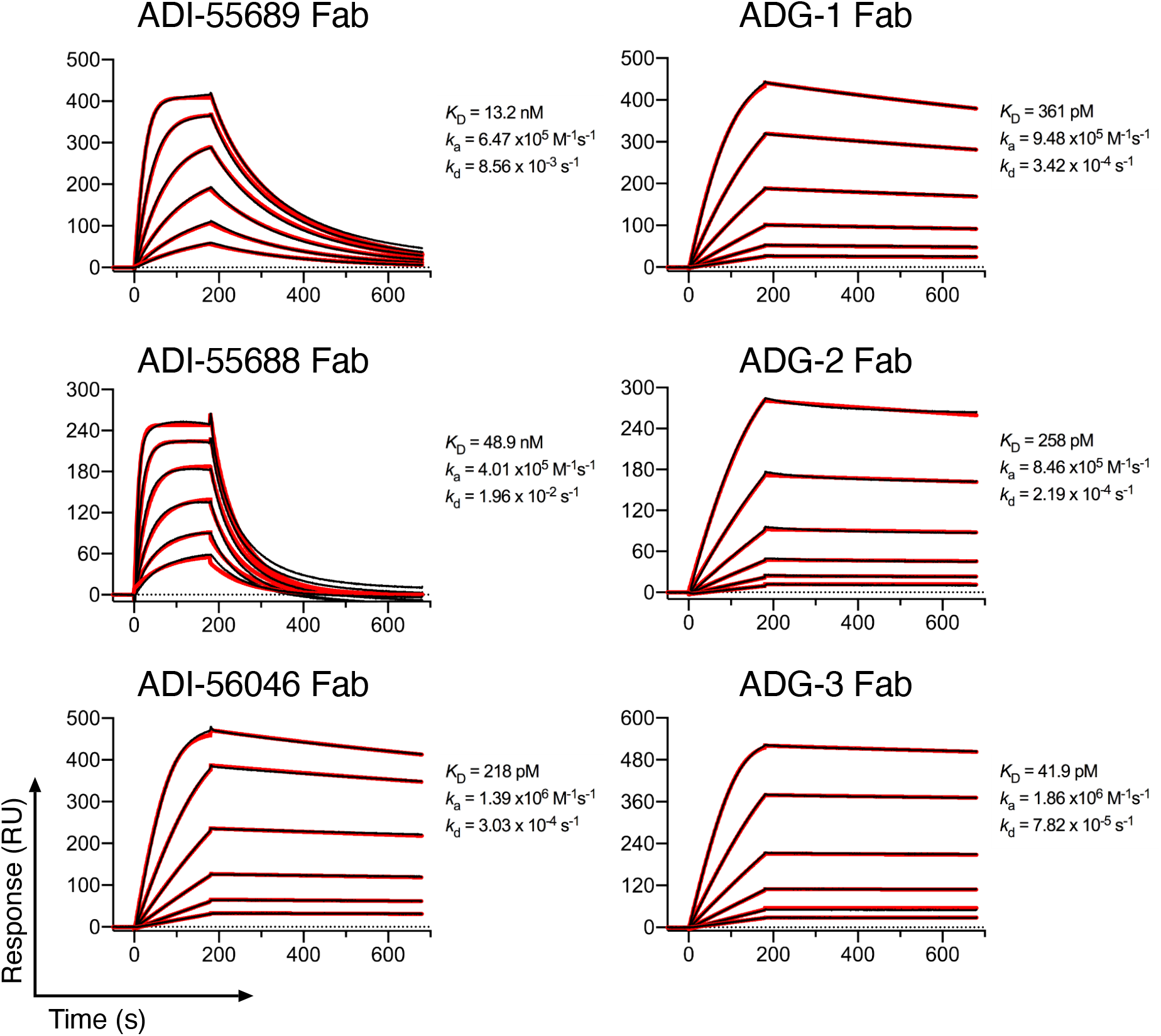
Binding kinetics of progenitor and affinity-matured Fabs. SPR sensorgrams showing binding of each Fab to the SARS-CoV-2 RBD-SD1 protein. Binding data are shown as black lines, and the best fits of a 1:1 binding model are shown as red lines.

**Figure S3.**
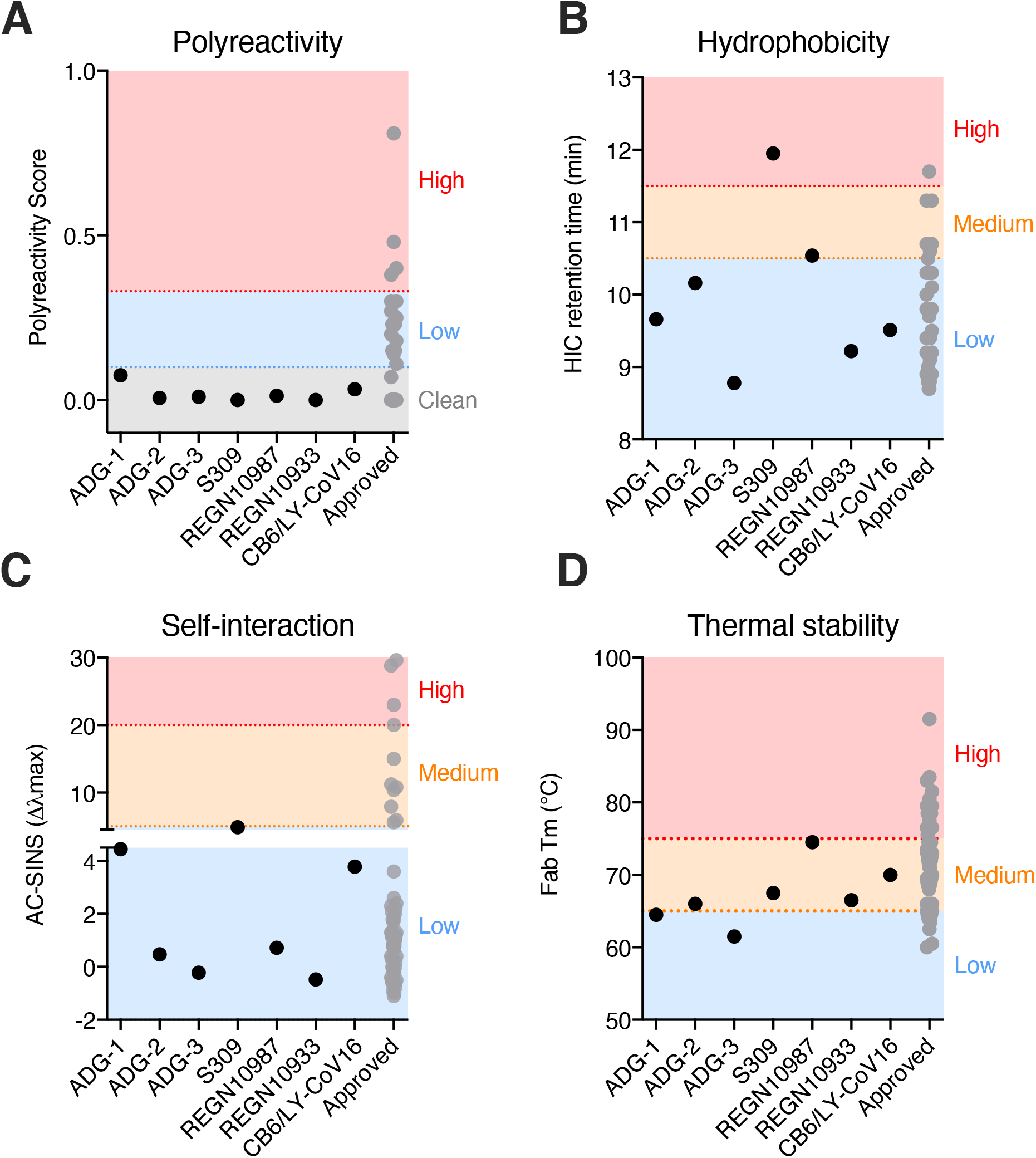
Biophysical properties of SARS-CoV-2 antibodies. **(A)** Antibody polyreactivity, as assessed based on binding to a previously described polyspecificity reagent (*37*). Binding was assessed by flow cytometry. The thresholds for high, low, and “clean” polyreactivity were defined based on a previously reported correlation between polyreactivity in this assay and serum half-life in humans (*22*). **(B)** Antibody hydrophobicity, as determined by hydrophobic interaction chromatography. **(C)** Antibody self-association propensity, as determined by affinity-capture self-interaction nanoparticle spectroscopy (AC-SINS). **(D)** Fab thermal stability, as determined by differential scanning fluorimetry (DSF). Forty-two clinically approved antibodies (*38*) were included in each assay as comparators and used to determine the thresholds for high/medium/low hydrophobicity, self-interaction propensity and thermal stability.

**Figure S4.**
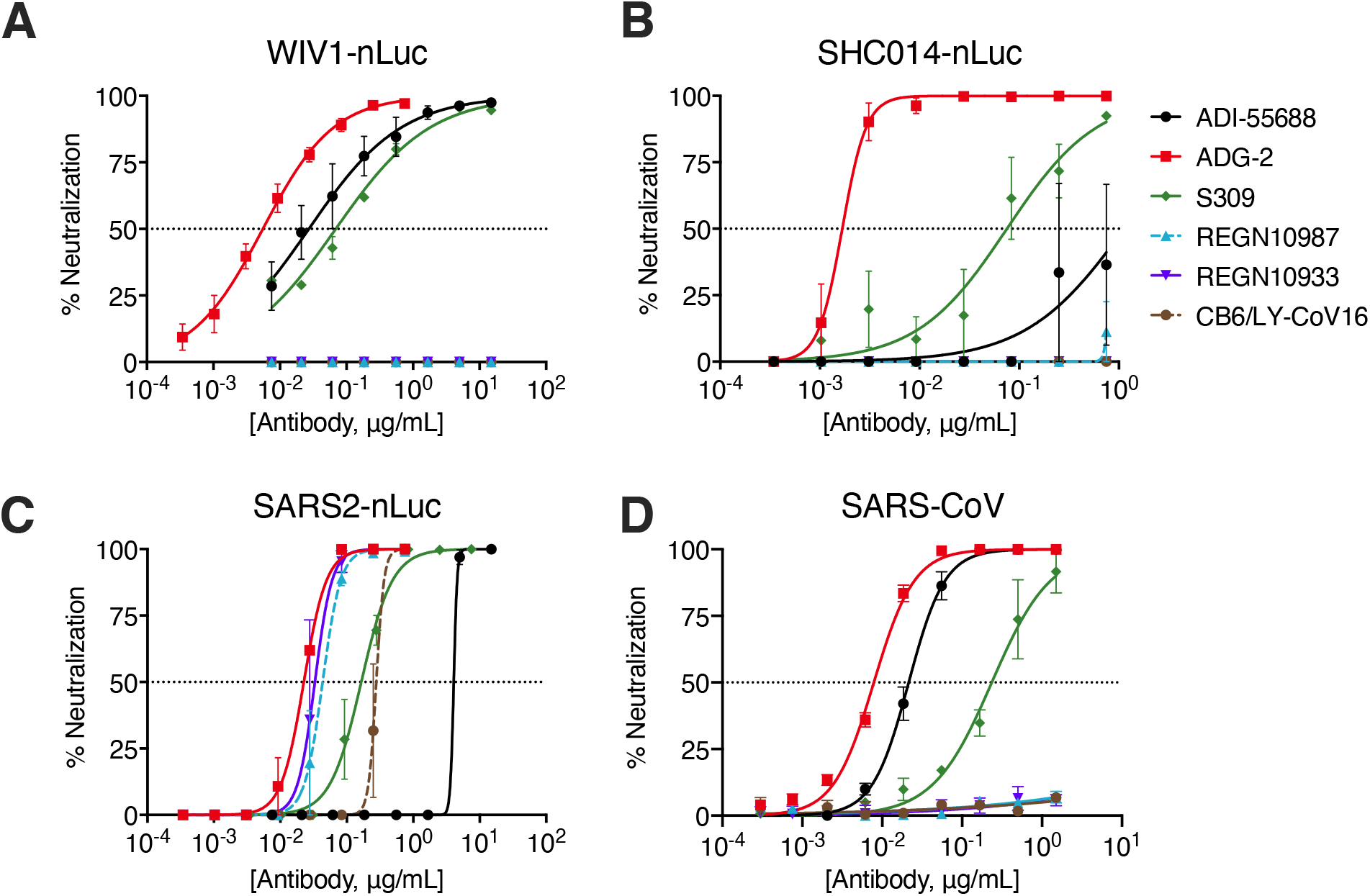
Representative neutralization curves for ADG-2 and SARS-CoV-2 clinical antibodies against authentic WIV-1 **(A)**, SHC014 **(B)**, SARS-CoV-2 **(C),** or SARS-CoV **(D)** on Vero target cells. Error bars represent standard deviations.

**Figure S5.**
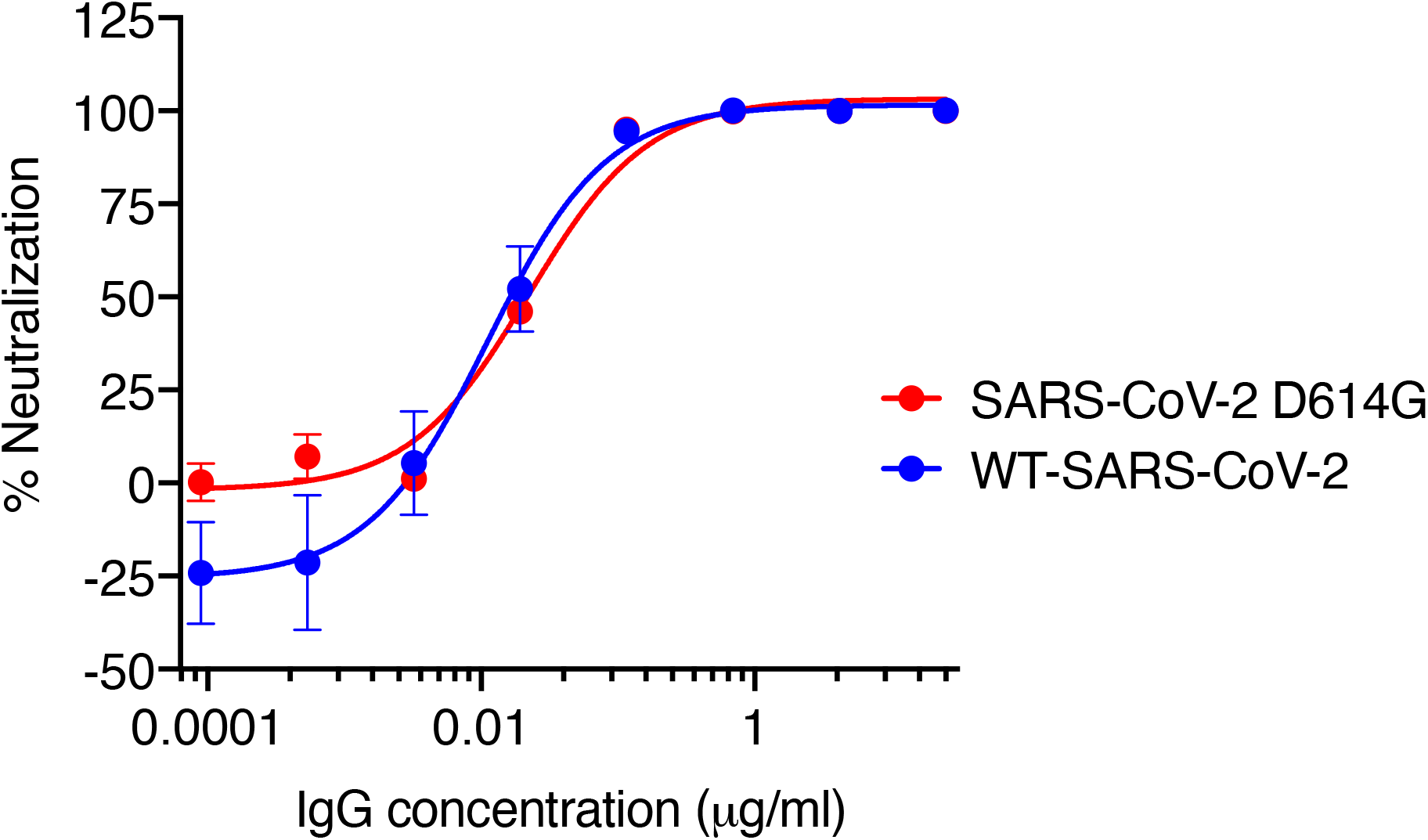
ADG-2 neutralization of SARS-CoV-2 D614G. Neutralizing activity of ADG-2 against WA1-SARS-CoV-2 and WA1-SARS-CoV-2 D614G viruses was assessed using a murine leukemia virus (MLV)-based pseudovirus assay and HeLa-hACE2 target cells.

**Figure S6.**
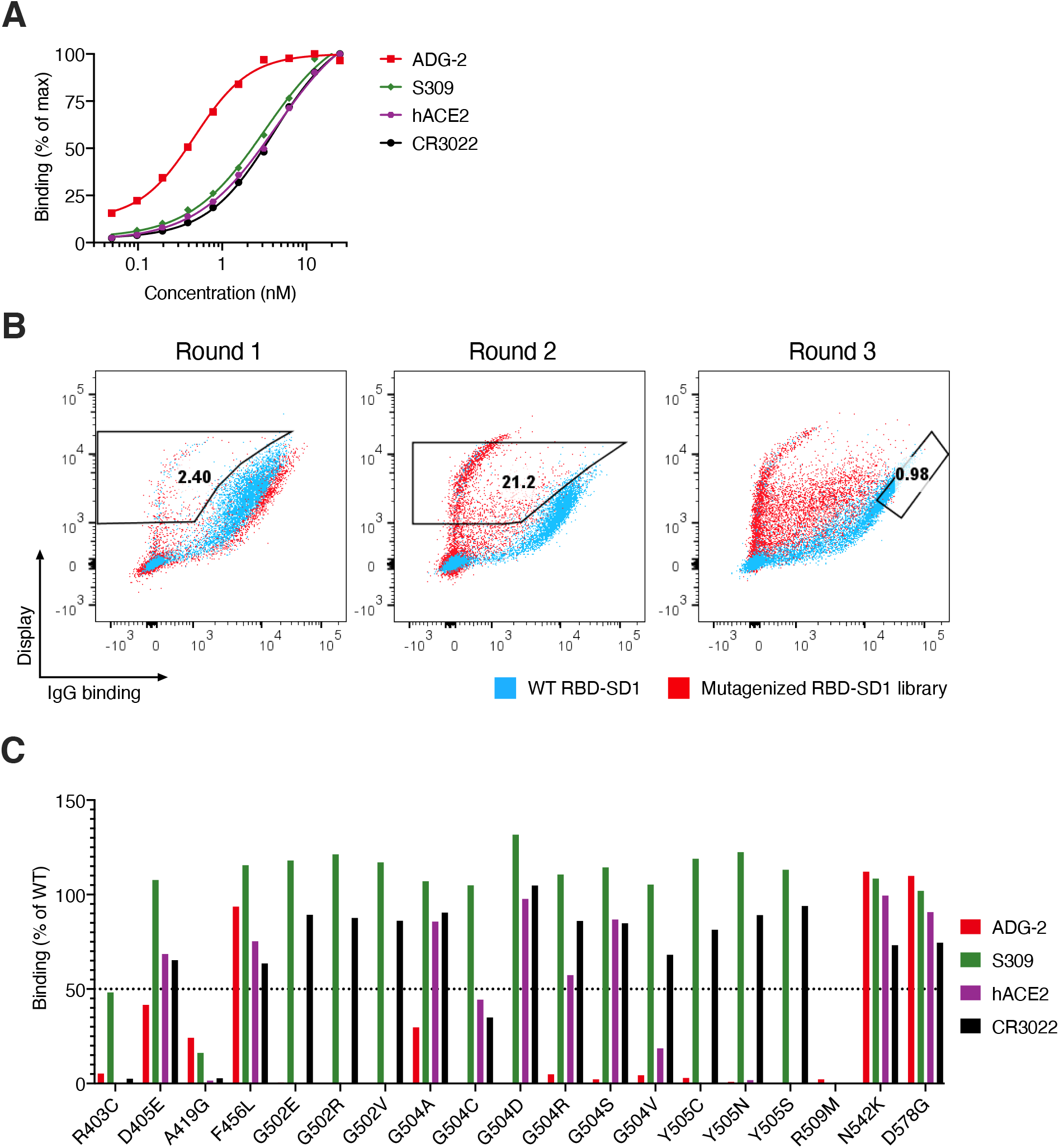
Fine epitope mapping using an RBD display library. **(A)** Binding titration curves of ADG-2, S309, CR3022, and recombinant hACE2-Fc protein on the WT yeast surface-displayed SARS-CoV-2 RBD, used to calculate K_D_^app^ and EC_80_ concentrations. **(B)** Flow cytometric selection of a mutagenized, yeast surface-displayed RBD library (red) to enrich for variants that display loss of binding to ADG-2 (round 1 and 2) but retain binding to antibodies (S309 and CR3022) that bind to epitopes distinct from the ADG-2 binding site (round 3). The round 1 and 2 gates indicate the yeast populations that were sorted for sequential rounds of selection and the round 3 gate indicates the yeast population that was sorted for individual colony isolation and sequencing. **(C)** Percent antibody or recombinant hACE2-Fc binding to yeast surface-displayed SARS-CoV-2 RBD variants relative to the WT SARS-CoV-2 RBD. Antibody binding was assessed at their respective EC_80_ concentrations for the WT RBD construct.

**Figure S7.**
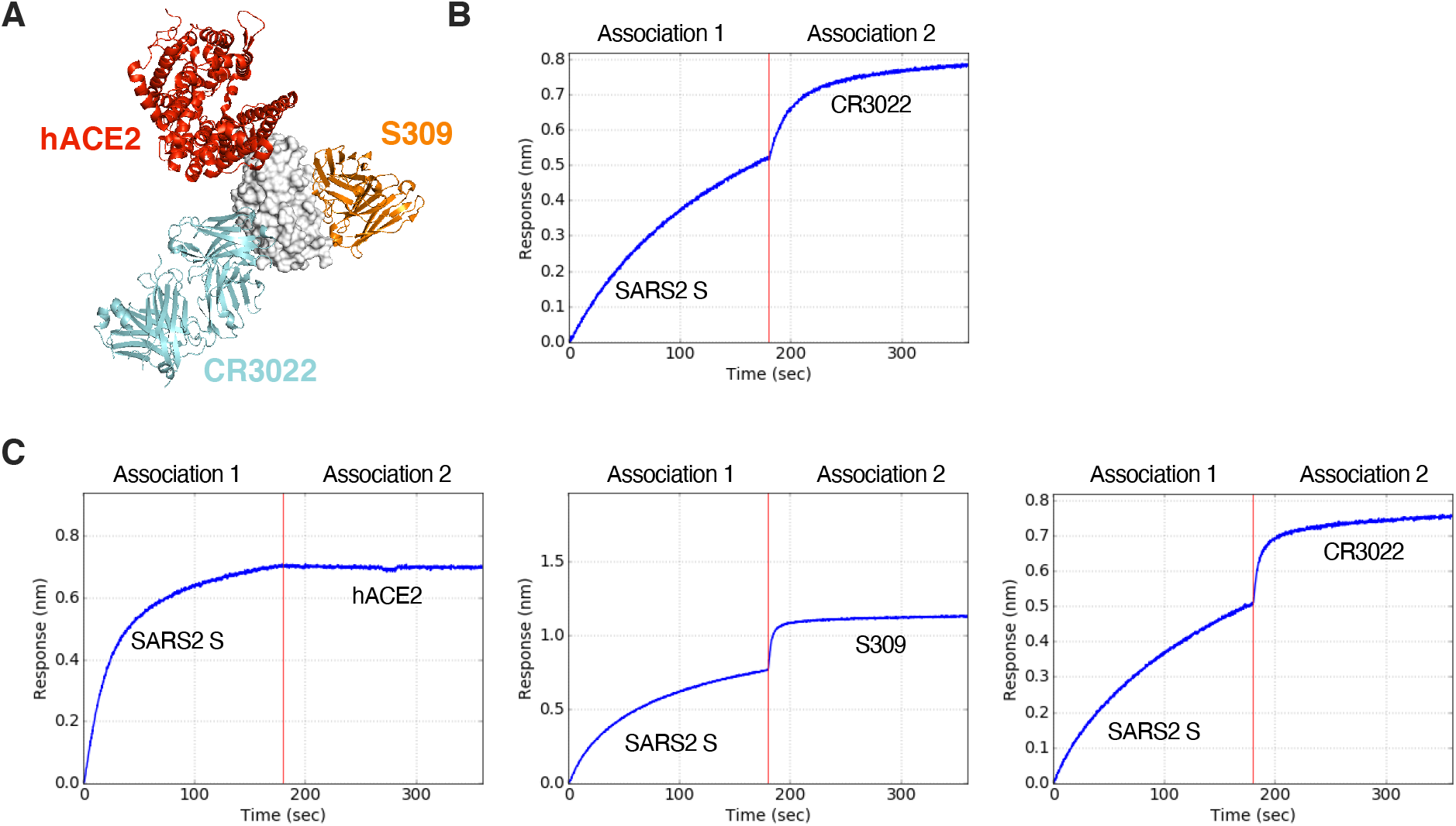
Competitive binding assays. **(A)** Structure of the SARS-CoV-2 RBD (white) bound to hACE2 (red) (PBD ID: 6M0J) docked with SARS-CoV-2 mAbs S309 (orange, PDB ID: 6WPS) and CR3022 (cyan, PBD ID: 6W41). **(B)** S309 and CR3022 sandwich binning sensorgram. The traces depict the association (0-180 seconds) of SARS-CoV-2 S to S309 captured on the probe followed by exposure (180-360 seconds) to CR3022. **(C)** ADG-2 and hACE2-Fc (left), S309 (middle), and CR3022 (right) sandwich binning sensorgrams. The traces depict the association (0-180 seconds) of SARS-CoV-2 S to ADG-2 captured on the probe followed by exposure (180-360 seconds) to hACE2-Fc, S309 IgG, or CR3022 IgG. Additional binding of the competitor protein indicates an unoccupied epitope (non-competitor), while no binding by the competitor protein indicates blocking (competitor) of the epitope by the IgG.

**Figure S8.**
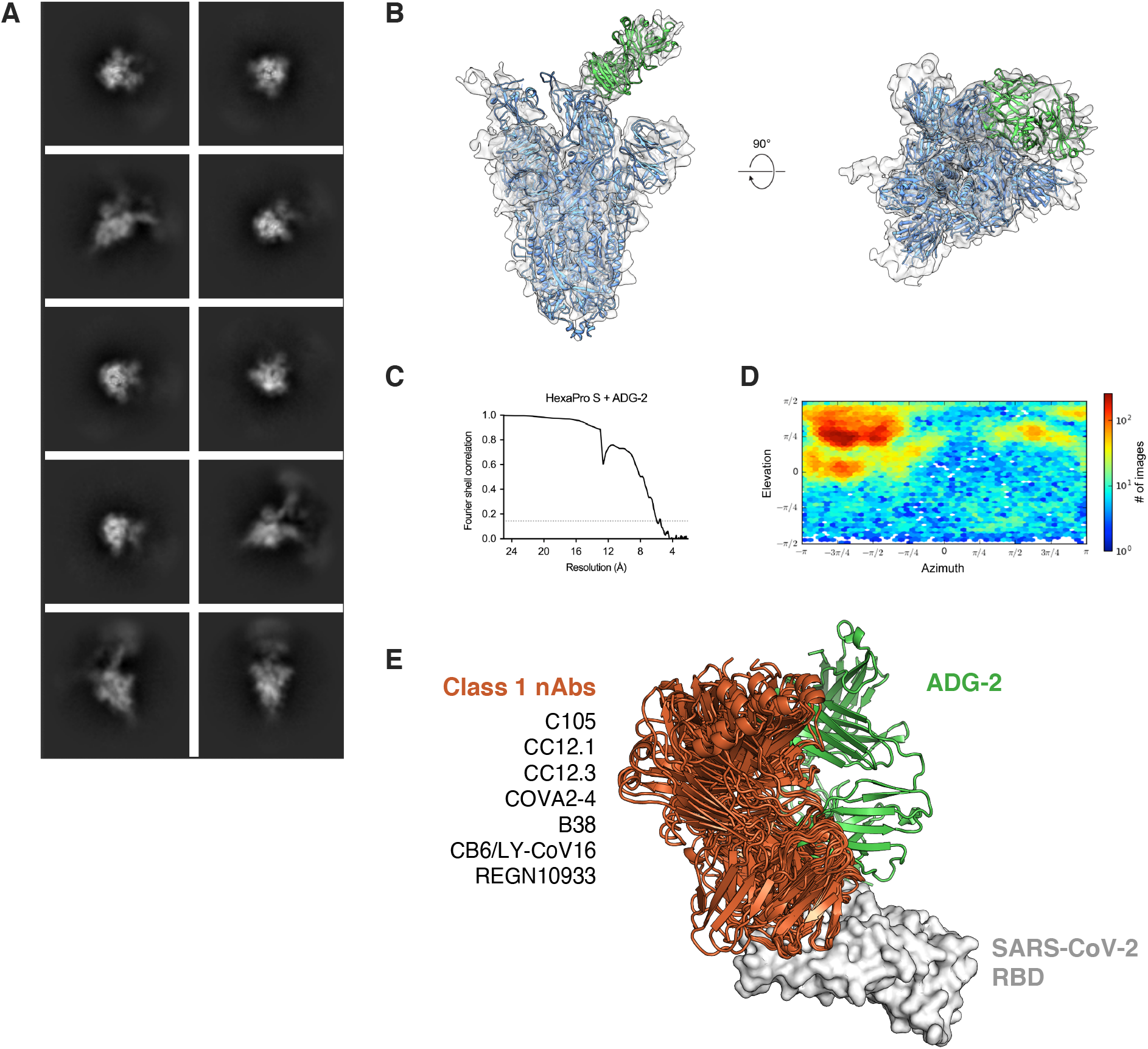
Cryo-EM validation and structural analysis of ADG-2 bound to the SARS-CoV-2 RBD. **(A)** Two-dimensional class averages of ADG-2 Fab bound to the SARS-CoV-2 spike. **(B)** Side and top views of the 5.94 Å cryo-EM reconstruction, with the map displayed as a transparent surface and high-resolution models of the SARS-CoV-2 spike (PDB ID: 6XKL) and a homologous Fab (PDB ID: 6APC) displayed as blue and green ribbons, respectively. **(C)** Fourier shell correlation (FSC) curve for the 3D reconstruction. The dashed line corresponds to an FSC value of 0.143. **(D)** The viewing distribution plot for the 3D reconstruction, calculated in cryoSPARC. **(E)** Cryo-EM reconstruction of the SARS-CoV-2 RBD (white) bound by ADG-2 Fab overlaid with high-resolution structures of several Class I SARS-CoV-2 neutralizing antibody Fabs (PDB IDs: 6XC4, 6XC2, 6XCN, 7JMO, 7BZ5, 6XDG and 7C01), shown as green and orange structures, respectively.

**Table S1.** Cryo-EM data collection and reconstruction statistics.

|  |  |
| --- | --- |
| Protein | SARS-CoV-2 HexaPro S + ADG-2 |
| EMDB | EMD-XXXXX |
| Microscope | FEI Titan Krios |
| Voltage (kV) | 300 |
| Detector | Gatan K3 |
| Magnification (nominal) | 22,500 |
| Pixel size (Å/pix) | 1.073 |
| Flux (e<sup>-</sup>/pix/sec) | 8.0 |
| Frames per exposure | 30 |
| Exposure (e<sup>-</sup>/Å<sup>2</sup>) | 36 |
| Defocus range (μm) | 0.6-3.5 |
| Micrographs collected | 4,748 |
| Particles extracted/final | 267,247/57,078 |
| Symmetry imposed | n/a (C1) |
| Map sharpening B-factor | 344.6 |
| Masked resolution at 0.143 FSC (Å) | 5.94 |

## Methods and Materials

### HeLa-hACE2 stable cell line

Stable human ACE2 (hACE2)-expressing HeLa cells for authentic SARS-CoV-2 neutralization assays were generated as previously described (*11*). Briefly, hACE2 (NM_001371415) was cloned into the pBOB vector and co-transfected with lentiviral vectors pMDL (Addgene #12251), pREV (Addgene #12253), and pVSV-G (Addgene #8454) into HEK293T cells using Lipofectamine 2000 (Thermo Fisher Scientific) according to the manufacturer’s protocol. Culture media was exchanged 16 hours post-transfection, and supernatant was harvested 32 hours post-transfection. Pre-seeded HeLa cells were transduced using harvested supernatant with 10 μg/mL polybrene (Sigma). At 12 hours post-transduction, cell surface expression of hACE2 was confirmed by flow cytometry.

### Authentic SARS-CoV neutralization assay

To generate authentic SARS-CoV, Vero African grivet monkey kidney cells (Vero E6, ATCC-CRL1586) were grown in Dulbecco’s Modified Eagle Medium (DMEM high glucose; Gibco, Cat # 11995065), 2% heat-inactivated fetal bovine serum (FBS, Atlanta Biologicals), 0.05% Trypsin-EDTA solution (Gibco), 1% Pen/Strep (Gibco), and 1% GlutaMAX (Gibco). Cells were infected with SARS-CoV/Urbani at a multiplicity of infection (MOI) of 0.01 and incubated at 37 °C with 5% CO_2_ and 80% relative humidity (RH). 50 hours post-infection, cells were frozen at −80 °C for 1 hour and then thawed at room temperature (RT). The supernatant was collected and clarified by centrifugation at 2500 × g for 10 minutes before aliquoting for storage at −80 °C. Virus neutralization was assessed as previously described (*11*). Briefly, SARS-CoV/Urbani (MOI = 0.2) was added to serial dilutions of antibodies and incubated for 1 hour at RT. The antibody-virus mixture was applied to monolayers of Vero E6 cells in a 96-well plate and incubated for 1 hour at 37 °C, 5% CO_2_ and 80% RH. Next, media was exchanged by washing cells once with 1x PBS and adding fresh cell culture media. At 24-hour post-infection, cells were washed out of media with 1x PBS to then be treated with formalin fixing solution, permeabilized with 0.2% Triton-X for 10 minutes at RT, and finally treated with blocking solution. Fixed and permeabilized cells were first stained with a primary antibody recognizing SARS-CoV nucleocapsid protein (Sino Biological), followed by secondary antibody staining with AlexaFluor 488-conjugated goat anti-rabbit antibody. Infected cells were enumerated by an Operetta high content imaging instrument, and data was analyzed using Harmony software (Perkin Elmer).

### MLV-SARS-CoV-2 pseudovirus neutralization assay

To generate the MLV pseudoviruses, pCDNA3.3 plasmids (ThermoFisher) encoding the WT (NC_045512) or D614G-variant SARS-CoV-2 spike gene with a 28 amino acid deletion at its C-terminus (IDT); a luciferase reporter gene plasmid (Addgene # 18760) modified with a cytomegalovirus (CMV) promoter to replace the internal ribosome entry site (IRES); and a murine leukemia virus (MLV) Gag-Pol plasmid (Addgene # 14887) were purified using the Endo-Free Plasmid Maxi Kit (Qiagen) according to manufacturer’s instructions. To generate single-round infection competent pseudoviruses, HEK293T cells were co-transfected with 2 μg of MLV Gag-Pol-, 2 μg of MLV luciferase-, and 0.5 μg of either SARS-CoV-2 WT S or SARS-CoV-2 D614G S-encoding plasmids in 6-well plates using Lipofectamine 2000 (Thermo FisherScientific), according to the manufacturer’s directions. Cell culture media was exchanged 16 hours post-transfection. At 48 hours post-transfection, the supernatant containing SARS-CoV-2 S-pseudotyped viral particles was harvested, aliquoted, and frozen at −80 °C.

Antibody neutralization of pseudoviruses was assessed via a luminescence-based assay on HeLa-hACE2 cells as previously described (*39*). The SARS-CoV-2 WT or D614G pseudovirus was mixed with serially diluted antibodies, incubated for 1 hour at 37 °C, and applied to 10,000 HeLa-hACE2 cells. Following infection for 42-48 hours at 37 °C, HeLa-hACE2 cells were lysed with 1x luciferase lysis buffer (25 mM Gly-Gly pH 7.8, 15 mM MgSO4, 4 mM EGTA, 1% Triton X-100). Luciferase intensity was measured on a luminometer using Bright-Glo luciferase substrate (Promega, PR-E2620) following manufacturer’s directions. Percentage of neutralization was calculated from sample and control relative units of light (RUL) according to the formula: 100*(1–[RUL_sample_ – RUL_backrground_]/[RUL_virus-only_ – RUL_background_]).

### Authentic WIV-1, SHC014 and SARS-CoV-2 nano-luciferase (nLuc) neutralization assays

Mouse-adapted SARS-CoV (MA15), mouse adapted SARS-CoV-2 (MA2) and WT SARS-CoV-2 nano-luciferase (nLuc) viruses were generated by CoV reverse genetics as described previously (*40*). WIV-1-nLuc and SHC014-nLuc were generated by replacing the CoV ORF7 and ORF8 regions with nano-luciferase. All nLuc viral assays were performed with Vero E6 cells, which were grown in DMEM high glucose media (Gibco, Cat # 11995065) supplemented with 10% fetal clone II (GE, Cat # SH3006603HI), 1% non-essential amino acids, and 1% Pen/Strep at 37°C and 5% CO_2_. Vero E6 cells were seeded at 2 × 10^4^ cells/well in a black-wall, tissue culture treated, 96-well plate (Corning, Cat # 3603) 24 hours prior to pseudovirus assays.

Antibodies were serially diluted in growth media and mixed at a 1:1 ratio with either 75 plaque forming units (PFU)/well SARS-CoV-MA15-nLuc, 100 PFU/well SARS-CoV-2-nLuc, 85 PFU/well SARS-CoV-2-MA2-nLuc, 20 PFU/well SHC014-nLuc, or 250 PFU/well WIV1-nLuc viruses and incubated at 37 °C for 1 hour. Virus and antibody mixture was then added to Vero E6 cells and incubated at 37 °C with 5% CO_2_ for 48 hours (SARS-CoV-MA15, SARS-CoV-2-MA2 and SARS-CoV-2-nLuc) or 24 hours (SHC014-nLuc and WIV1-nLuc). Luciferase activities were measured by the Nano-Glo Luciferase Assay System (Promega Cat. #N1130), following the manufacturer’s protocol, using a SpectraMax M3 luminometer (Molecular Devices). Percent inhibition was calculated by the following equation: 100*(1 – [RLU_sample_/ RLU_mock-treatment_]). Half-maximal inhibitory concentrations (IC_50_) were calculated for each condition by curve-fitting with non-linear regression.

### Authentic SARS-CoV-2 neutralization assay

Authentic SARS-CoV-2 virus was produced in Vero E6 cells as described previously (*23*). Vero E6 cells were grown overnight in complete DMEM (Corning, Cat # 15-013-CV) supplemented with 10% FBS, 1x Pen/Strep (Corning, Cat # C20-002-CL), and 2 mM L-Glutamine (Corning, Cat # 25-005-CL) at 37°C and 5% CO_2_. Cells were incubated with 2 mL of SARS-CoV-2 strain USA-WA1/2020 (BEI Resources, Cat # NR-52281) at MOI of 0.5 for 30 minutes at 34°C and 5% CO_2_, followed by direct addition of 30 mL of complete DMEM. At 5 days post-infection, the supernatant was collected and centrifuged at 1000 × g for 5 minutes, passed through 0.22 μM filters, and frozen at −80 °C for future use.

Antibody neutralization against authentic SARS-CoV-2 was assessed using both Vero E6 and HeLa-hACE2 target cells. Both types of target cells were grown in complete DMEM at 37°C and 5% CO_2_. For neutralization assays, HeLa-hACE2 or Vero E6 target cells were seeded in a 96-well half-well plate at approximately 8000 cells/well suspended in 50 μL complete DMEM and grown overnight. 1,000 plaque forming units (PFU)/well of SARS-CoV-2 was added to titrating amounts of antibody and incubated for 30 minutes. The virus-antibody mixture was subsequently incubated with either HeLa-hACE2 or Vero E6 cells for 24 hours at 37°C and 5% CO_2_. Following incubation, the infection media was removed. Cells were submerged in 4% formaldehyde for 1 hour, followed by three cycles of washing with PBS, and incubated with 100 μL/well of permeabilization buffer (1x PBS with 1% Triton-X) with gentle shaking. The plates were then blocked with 100 μL of 3% [w/v] bovine serum albumin (BSA) for 2 hours at room temperature (RT) and subsequently washed out of blocking solution with wash buffer (1x PBS with 0.1% Tween-20).

SARS-CoV-2 viruses were detected with a mixture of CC6.29, CC6.33, L25-dP06E11, CC12.23, and CC12.25 antibodies, previously derived from a cohort of convalescent SARS-CoV-2 donors (*23*). Pooled antibodies were added to wells at a concentration of 2 μg/mL (50 μL/well) and incubated for 2 hours at RT. Cells were subsequently washed 3 times with wash buffer, stained with 0.5 μg/mL peroxidase-conjugated AffiniPure goat anti-human IgG (Jackson ImmunoResearch Laboratories, Inc, Cat # 109-035-088) for 2 hours at RT, and followed by 6 washes with wash buffer. Freshly prepared HRP substrate (Roche, Ca # 11582950001) at a 100:1 volume ratio of Solution A:B was added to each well. Chemiluminescence was measured using a microplate luminescence reader (BioTek, Synergy 2).

A standard curve of serially diluted virus from 3000 to 1 PFU was plotted against relative light units (RLU) using a 4-parameter logistic regression as follows: *y = a + (b – a) / (1 + (x / x_0_)^c^),* where *y* = variable in RLU, *x* = variable in PFU and *a*, *b*, *c* and *x*0 are parameters fit by the standard curve. Using parameters generated by the standard curve, sample RLU values were converted into PFU values (*x* = *x*0 × log_c_ [(*b – y*) / (*y – a*)]), and percentage neutralization was calculated with the following equation: % Neutralization = 100 × [(VC – ADG-2 treated) / (VC – CC)], where VC = average of vehicle-treated control and CC = average of cell only control, both variables in PFU values. Half maximal inhibitory concentration (IC_50_) values were determined using curve fitting using non-linear regression.

### Mammalian expression and purification of recombinant SARS-CoV-2 S antigens

Plasmids encoding residues 319–591 of SARS-CoV-2 S with a C-terminal monomeric human IgG Fc-tag and an 8x HisTag (SARS-CoV-2 RBD-SD1); residues 1–1208 of the SARS-CoV-2 spike with a mutated S1/S2 cleavage site, proline substitutions at positions 986 and 987, and a C-terminal T4-fibritin trimerization motif, an 8x HisTag, and a TwinStrepTag (SARS-CoV-2 S-2P); or residues 1–1208 of the SARS-CoV-2 spike with a mutated S1/S2 cleavage site, proline substitutions at positions 817, 892, 899, 942, 986, and 987, a C-terminal T4-fibritin trimerization motif, an 8x HisTag, and a TwinStrepTag (SARS-CoV-2 HexaPro S) were transiently transfected into FreeStyle293F cells (Thermo Fisher) using polyethylenimine. Two hours post-transfection, cells were treated with kifunensine to ensure uniform, high-mannose glycosylation. Cell supernatants were harvested after 6 days of protein expression. SARS-CoV-2 RBD-SD1 was purified using Protein A resin (Pierce) and SARS-CoV-2 S-2P and SARS-Cov-2 HexaPro S were purified using StrepTactin resin (IBA). Affinity-purified SARS-CoV-2 RBD-SD1 was further purified over a Superdex75 column (GE Life Sciences). SARS-CoV-2 S-2P and SARS-CoV-2 HexaPro S were purified over a Superose6 Increase column (GE Life Sciences).

### In vitro affinity maturation of ADI-55688, ADI-55689, and ADI-56046

For each antibody, the complementarity-determining regions (CDRs) 1, 2, and 3 of the heavy- and light-chains were diversified separately via oligo-based mutagenesis using NNK-randomized oligos spanning CDRH1, CDRH2, CDRH3, CDRL1, CDRL2, and CDRL3 (IDT). Overlap-extension PCR was used to assemble and amplify forward-priming NNK oligos and reverse-priming oligo pools covering framework regions 1-4 with added homology to the CDR oligos to generate full-length variable regions. For the CDRH1/CDRH2/CDRH3 selections, heavy-chain variable regions (HCFR1-HCFR4) and the unmutated light-chain variable regions of the nominated parent were recombined *in situ* by homologous recombination with linearized vector to create a yeast library of 1 × 10^7^ diversity via electroporation.

Heavy- and light-chain libraries of each parent antibody (ADI-55688, ADI-55689, and ADI-56046) were subject to two rounds of selection for binding to a recombinant SARS-CoV-2 S1 protein (Sino Biological, Cat # 40591-V08H). Induced yeast libraries covering at least 10-fold of their respective diversities were incubated with 10 or 1 nM biotinylated SARS-CoV-2 S1 protein under equilibrium conditions. Yeast was washed twice in PBSF (1x PBS, 0.1% [w/v] BSA), stained with anti-human LC-FITC (Southern Biotech), Streptavidin 633 (Invitrogen, Cat # S21375), and propidium iodide (Invitrogen, Cat # P1304MP) for 15 minutes on ice. Labeled cells were subsequently washed twice and resuspended in PBSF before sorting on a BD FACS Aria II (Becton Dickerson). Gates were drawn for cells with improved S1 binding over parental clones.

Following two rounds of sorting, the variable heavy and light regions of enriched output clones were recombined to generate new CDRH1/CDRH2/CDRH3/CDRL1/CDRL2/CDRL3 libraries. An additional two rounds of selections were performed as described above. Sorted yeast from the final round of selection were resuspended in SDCAA media and plated on SDCAA agar plates for single colony isolation and sequencing.

### Expression and purification of IgGs and Fab fragments

Monoclonal antibodies ADI-55688, ADI-55689, and ADI-56046, as well as their progeny, were produced as full-length IgG_1_ proteins in *S. cerevisiae* cultures, as previously described (*11*). Briefly, yeast cultures were incubated in 24-well plates placed in Infors Multitron shaking incubators at 30 °C, 650 rpm, and 80% relative humidity. After 6 days, the supernatants containing the IgGs were harvested by centrifugation and purified by protein A-affinity chromatography. The bound IgGs were eluted with 200 mM acetic acid with 50 mM NaCl (pH 3.5) into 1/8 [v/v] 2 M HEPES (pH 8.0) and buffer-exchanged into PBS (pH 7.0).

ADG1-3 and benchmark SARS-CoV-2 mAbs REGN10933, REGN10987, CB6/LY-CoV016, and S309 were expressed in CHO cells as full-length IgG_1_ proteins. The VH- and VL-encoding gene fragments were subcloned into heavy- and light-chain vectors and transiently transfected into CHO cells. After 6 days, the supernatants containing the IgGs were harvested by centrifugation and purified by protein A-affinity chromatography. Bound IgGs were eluted and further purified by size exclusion chromatography (SEC) to at least 95% purity, then buffer-exchanged into 150 mM NaCl with 20 mM histidine, pH 6.0.

Fab fragments for structural studies were generated by digestion with papain for 2 hours at 30 °C, followed by the addition of iodoacetamide to terminate the reaction. To remove the Fc fragments and any undigested IgG fractions, the mixtures were passed over Protein A agarose. The flow-through of the Protein A resin was then passed over CaptureSelect™ IgG-CH1 affinity resin (ThermoFisher Scientific) and the captured Fabs were eluted with 200 mM acetic acid with 50 mM NaCl (pH 3.5) into 1/8 [v/v] 2 M HEPES (pH 8.0), followed by buffer exchange into PBS (pH 7.0).

### Surface plasmon resonance Fab kinetic binding measurements

SEC-purified SARS-CoV-2 RBD-SD1 was immobilized to a Ni-NTA sensor chip in a Biacore X100 (GE Life Sciences) to a response level of ~500 RUs. Fabs were then injected at increasing concentrations, ranging from 18.75-300 nM (ADI-55688), 1.56-25 nM (ADI-56046), 6.25-100 nM (ADI-55689), or 1.25-20 nM (ADG-1, ADG-2, ADG-1). The sensor chip was doubly regenerated between cycles using 0.35 M EDTA and 0.1 M NaOH. The resulting data were double-reference subtracted and fit to a 1:1 binding model using Biacore Evaluation Software.

### Competitive binding experiments using biolayer-interferometry

Competition of ADG-2 with recombinant hACE2-Fc protein (Sino Biological, Cat # 10108-H02H). CR3022, and S309 for binding to soluble SARS-CoV-2 S trimer was assessed using the ForteBio Octet HTX (Sartorius Bioanalytical Instruments). All reagents were diluted to 100 nM in PBSF. Anti-heavy-chain (AHC) sensor tips were loaded with S309 or ADG-2 IgG, followed by exposure to an inert IgG to block any remaining Fc capture sites. Tips were subsequently equilibrated in PBSF for 30 minutes. IgG-loaded sensor tips were transferred to wells containing hACE2, CR3022, or S309 to check for any interaction with the loaded IgG. Sensor tips were then loaded in wells containing fresh PBSF buffer (60 seconds), followed by exposure to SARS-CoV-2 S protein (180 seconds), and lastly, exposure to hACE2, CR3022, or S309 (180 seconds). Data were cropped to include only SARS-CoV-2 S protein and hACE2, CR3022, or S309 exposure steps and aligned by x- and y-axes using ForteBio Data Analysis software version 11.1.3.10.

### Antibody-dependent natural killer cell activation and degranulation (ADNKDA)

Primary human NK cells were enriched from the peripheral blood of human donors using RosetteSep Human NK cell Enrichment Cocktail (Stem Cell Technologies, Cat #15065) and cultured overnight in RPMI-1640 (Corning, Cat # 15-040-CV) supplemented with 10% FBS (Hyclone, Cat # SH30071.03), 1% Pen/Strep (Gibco, Cat # 15070-063), 1% L-Glutamine (Corning, Cat # 25-005-CI), 1% HEPES (Corning, Cat # 25-060-CI) and 5 ng/mL recombinant human IL-15 (StemCell Technologies, Cat # 78031). Recombinant SARS-CoV-2 receptor binding domain was coated onto MaxiSorp 96-well plates (Thermo Scientific, Cat # 442404) at 200 ng/well at 4 °C overnight. Wells were washed with PBS and blocked with 5% BSA prior to addition of antibodies that were diluted in a five-fold dilution series in PBS (10 μg/mL - 0.32 ng/mL) and incubation for 2 h at 37 °C. Unbound antibodies were removed by washing with PBS were added at 5 × 10^4^ cells/well in the presence of 4 μg/mL brefeldin A (Biolegend, Cat # 420601), 5 μg/mL GolgiStop (BD Biosciences, Cat # 554724) and anti-CD107a antibody (Clone H4A3 PE-Cy7, Biolegend, Cat # 328618) for 5 hours. Cells were stained for surface expression of CD16 (Clone 3G8 Pacific Blue, Biolegend, Cat # 302032), CD56 (clone 5.1H11 AlexaFluor488, Biolegend, Cat # 362518) and CD3 (clone UCHT1 Alexa Fluor700, Biolegend, Cat # 300424). Cells were fixed and permeabilized with Fix/Perm (Biolegend, Cat # 421002) according to the manufacturer’s instructions to stain for intracellular IFNγ (Clone B27 PE, Biolegend, Cat # 506507) and TNFα (clone Mab11 APC, Biolegend, Cat # 502912). Cells were analyzed on a Cytek Aurora spectral flow cytometer.

### Antibody-dependent cellular phagocytosis (ADCP) with monocytes and neutrophils

For ADCP assays with neutrophils, HL-60 promyeloblast cells (ATCC, Cat # CCL-240) were maintained in Iscove’s Modified Dulbecco’s Medium (ATCC, Cat # 30-2005) with 20% fetal bovine serum and 1% Pen/Strep. HL-60 cells were differentiated into neutrophils by growth for 5 days in the presence of 1.3% DMSO. Recombinant SARS-CoV-2 RBD protein was coupled to fluorescent beads (Thermo Scientific, Cat # F8819) by carbodiimide coupling. Antibodies were diluted in a five-fold dilution curve in HL-60 culture medium (1000 - 0.32 ng/mL) and incubated with RBD-coated beads for 2 hours at 37 °C. Cells (5 × 10^4^/well) were incubated for 18 hours at 37 °C. Cells were then stained for CD11b (Clone M1/70 APC-Fire750, Biolegend, Cat # 101262) and CD16 (Clone 3G8 Pacific Blue, Biolegend, Cat # 302032), fixed with 4% paraformaldehyde, and analyzed by flow cytometry. CD11b+ and CD16+ cells were analyzed for uptake of fluorescent beads. A phagocytic score was determined using the following formula: (percentage of FITC^+^ cells)*(geometric mean fluorescent intensity (gMFI) of the FITC^+^ cells)/100,000.

For ADCP assays with monocytes, THP-1 monocytes were maintained in RPMI-1640 supplemented with 10% FBS, 1% Pen/Strep, 1% L-glutamine, and β-mercaptoethanol. Recombinant SARS-CoV-2 RBD-coated beads were generated as described above. Antibodies were diluted in a five-fold dilution curve in THP-1 culture medium to (5000–0.064 ng/mL) and incubated with RBD-coated beads for 2 h at 37 °C. Unbound antibodies were removed by centrifugation prior to the addition of THP-1 cells at 2.5 × 10^4^ cells/well. Cells were fixed with 4% paraformaldehyde and analyzed by flow cytometry. A phagocytic score was determined as described above.

### Antibody-mediated complement deposition (ADCD)

Recombinant SARS-CoV-2 receptor binding domain-coated beads were generated as described for ADCP assays. Antibodies were diluted in a five-fold dilution series in RPMI-1640 (5000 - 0.064 ng/mL) and incubated with RBD-coated beads for 2 hours at 37 °C. Unbound antibodies were removed by centrifugation prior to the addition of reconstituted guinea pig complement (Cedarlane Labs, Cat # CL4051) and diluted in veronal buffer supplemented with calcium and magnesium (Boston Bioproducts, Cat # IBB-300) for 20 minutes at 37 °C. Beads were washed with PBS containing 15 mM EDTA, and stained with an FITC-conjugated anti-guinea pig C3 antibody (MP Biomedicals, Cat # 855385). C3 deposition onto beads was analyzed by flow cytometry. The gMFI of FITC for all beads was measured.

### Polyreactivity assay

Polyspecificity reagent binding of antibodies was performed as described previously (*37*). Briefly, soluble membrane protein (SMP) and soluble cytosolic protein (SCP) fractions were extracted from Chinese hamster ovary (CHO) cells and biotinylated using NHS-LC-Biotin (Thermo Fisher Scientific) reagent. Yeast-presented IgGs were incubated with 1:10 diluted stock of biotinylated SMP and SCP for 20 minutes on ice, followed by two washes with PBSF, and stained with 50 μL of a secondary labeling mix containing ExtrAvidin-R-PE (Sigma-Aldrich), anti-human LC-FITC (Southern Biotech), and propidium iodide (Invitrogen) for 15 minutes on ice. Cells were subsequently washed with PBSF and resuspended in PBSF for flow cytometric analysis on a BD FACS Canto II (BD Biosciences). Polyreactivity scores were also reported for 42 previously described clinical antibodies for comparison (*38*).

### Affinity-capture self-interaction nanoparticle spectroscopy (AC-SINS)

To measure the propensity for antibodies to self-associate, AC-SINS was performed as previously described (*41*). Briefly, polyclonal goat anti-human IgG Fc antibodies (capture; Jackson ImmunoResearch Laboratories) and polyclonal goat non-specific antibodies (non-capture; Jackson ImmunoResearch Laboratories) were buffer exchanged into 20 mM sodium acetate (pH 4.3) and concentrated to 0.4 mg/mL. A 4:1 volume ratio of capture:non-capture was prepared and further incubated at a 1:9 volume ratio with 20 nm gold nanoparticles (AuNP; Ted Pella Inc.) for 1 hour at room temperature (RT). Thiolated PEG (Sigma-Aldrich) was then used to block empty sites on the AuNP and filtered via a 0.22 μm PVDF membrane (Millipore). Coated particles were subsequently added to the test antibody solution and incubated for 2 hours at RT before measuring absorbance from 510 to 570 nm on a plate reader. Data points were fit with a second-order polynomial in Excel to obtain wavelengths at maximum absorbance. Values are reported as the difference between plasmon wavelengths of the sample and background (Δλ_max_). AC-SINS values were also reported for 42 previously described clinical antibodies for comparison (*38*).

### Fab thermal stability

Apparent melting temperatures (T_m_^App^) of Fab fragments were obtained as previously described (*42*). Briefly, 20 μl of test antibody solution at 1 mg/mL was mixed with 10 μl of 20 × SYPRO orange. The plate was scanned with a CFX96 Real-Time System (BioRad) from 40 °C to 95 °C at a rate of 0.25 °C/minute. T_m_^App^ was calculated from the primary derivative of the raw data via the BioRad analysis software. Melting temperatures were also reported for 42 previously described clinical antibodies for comparison (*38*).

### Hydrophobic interaction chromatography (HIC)

Antibody hydrophobicity was evaluated using HIC as previously described (*43*). Briefly, test antibody samples were diluted in phase A solution (1.8 M ammonium sulfate and 0.1 M pH 6.5 sodium phosphate) to a final concentration of 1.0 M ammonium sulfate. A linear gradient from phase A solution to phase B solution (0.1 M pH 6.5 sodium phosphate) was run for 20 minutes at a flow rate of 1.0 mL/minute using the Sepax Proteomix HIC butyl-NP5 column. Peak retention times were obtained from monitoring UV absorbance at 280 nm. Hydrophobicity values were also reported for 42 previously described clinical antibodies for comparison (*38*).

### Sarbecovirus phylogeny and alignment

Representative sarbecovirus RBD-SD1 sequences were selected based on previously curated sequence sets (*25, 26*). Four additional ACE2-utilizing clade I sarbecoviruses (Frankfurt 1, CS24, Civet 007-2004, and A021) not represented in these curated sets were included for added diversity at the RBD-ACE2 interface. A limited set of clade 2 and clade 3 viruses, which do not utilize ACE2 as a target receptor (*26*), were included as controls. A phylogram of sarbecoviruses was generated using maximum likelihood analysis of MAFFT-aligned RBD-SD1 sequences. Multiple sequence alignment of sarbecovirus RBD sequences was visualized in Jalview. Amino acid sequences for each sarbecovirus were colored by percentage sequence identity and the overall degree of conservation per residue was calculated as a numerical index weighted by physio-chemical properties of amino acids (*44*).

### GISAID analysis of circulating SARS-CoV-2 variants

Genome sequences were downloaded from the GISAID database (*28*) and aligned pairwise against the reference Wuhan-Hu-1 sequence (ENA QHD43416.1) via an internal implementation of the Needleman–Wunsch algorithm to extract all RBD-SD1 sequences using amino acid residues 319 to 591 of the Wuhan-Hu-1 spike sequence. Incomplete RBD-SD1 nucleotide sequences and those containing ambiguous (“n”) base calls, plus translated sequences including “X”, “*”, or “-,” were excluded from further analysis. RBD-SD1 sequence variants observed at least 6 times out of 63551 sequences analyzed as of July 14, 2020, as well as several literature controls and antibody escape mutants (*24, 27*) observed in the GISAID database, were compiled as a panel 36 variants to assess antibody binding. Sequence frequencies were updated October 19, 2020 and used to calculate each percent prevalence.

### Cloning and expression of SARS-CoV-2 variant and homologous sarbecovirus RBD constructs

The spike RBD-SD1 of SARS-CoV-2 (residues 319 to 591, as defined by Uniprot: P0DTC2) and additional related sarbecoviruses (HKU3, ENA AAY88866.1; Rf1-2004, ENA ABD75323.1; BM48-31, ENA ADK66841; Pangolin_GX-P2V GISAID MT072864.1; RaTG13, ENA QHR63300.2; SARS-CoV-2, ENA QHD43416.1; GD-Pangolin, ENA MT121216.1; Rs4231, ENA ATO98157.1; WIV1, ENA AGZ48831.1; Civet 007-2004, ENA AAU04646.1; A021, ENA AAV97986.1; Frankfurt 1, ENA BAE93401.1; SARS-CoV-1, ENA AAP13441; CS24, ENA ABF68959; LYRa11, ENA AHX37558.1; Rs4081, ENA KY417143.1) were obtained as gBlocks (IDT) and cloned into a yeast surface-display expression vector encoding a flexible Gly4Ser linker and a hemagglutinin (HA) epitope tag at its N-terminus. Two consecutive Gly4Ser linkers connect RBD-SD1 to Aga2p at the C-terminus. Circulating SARS-CoV-2 variant sequences (described above) were cloned into the same expression vector. The A352S variant was excluded due to an error present in the provided gBlock. Plasmids were transformed into *S. cerevisiae* (EBY100) using the Frozen-EZ Yeast Transformation II Kit (Zymo Research) following the manufacturer’s protocol and recovered in selective SDCAA media.

For induction of RBD expression, fresh yeast cultures were inoculated at 0.2 OD_600_ in selective SDCAA media and grown at 30 °C and 180 rpm until cultures reached an 0.8-1.0 OD_600_. Cells were centrifuged at 2,400 × g for 3 minutes, resuspended in an equal volume of SGCAA (6.7 g/L Yeast Nitrogen Base, 4.0 g/L drop out amino acid mix, 0.46 g/L NaH_2_PO_4_, 0.88 g/L Na_2_HPO_4_, 7.7 g/L NaCl, 2% galactose, 2% raffinose), and incubated for 16 to 20 hours at 20 °C and 200 rpm.

### Antibody binding to yeast surface-displayed RBD variants

To assess binding breadth, IgGs and recombinant hACE2 (expressed in a bivalent format as a C-terminal IgG1 Fc conjugate; Sino Biological, Cat # 10108-H02H) were tested against the panel of 17 sarbecovirus RBDs. Initially, binding was determined at a single 100 nM concentration of IgG or hACE2. Induced cells (0.2 OD_600_ / well) were aliquoted into 96-well plates and washed out of SGCAA media with PBSF. Cells were resuspended in 100 μL of 100 nM IgG or hACE2 and incubated at room temperature for 30 minutes. Cells were subsequently washed twice with PBSF and labeled with 50 μL of APC-conjugated monoclonal mouse anti-hemagglutinin tag (HA).11 antibody (BioLegend, Cat # 901524), PE-conjugated goat anti-human IgG polyclonal antibodies (Southern Biotech, Cat # 2040-09), and propidium iodide (Invitrogen, Cat # P1304MP) for 20 minutes on ice. For each sarbecovirus RBD, a secondary reagent control was included. Cells were washed twice with PBSF before analyzing via flow cytometry on a BD FACS Canto II (BD Biosciences).

To account for differences in RBD expression across sarbecoviruses, binding signal was normalized to HA-tag signals (MFI_anti-human IgG PE_/MFI_anti-HA APC_). Binding with normalized ratios below 1.0 were considered non-binding (NB) at the concentration tested. Those with ratios above 1.0 were titrated between 100 nM to 0.048 nM to calculate their apparent binding affinity (*K*DApp). Mean anti-human IgG PE MFI signal was normalized according to the formula: (MFI_sample_ – MFI_minimum_)*100/(1 – MFI_minimum_) and fitted as nonlinear regression curves in GraphPad Prism using the following equation: Y=Y_x=minimum_ + X*(Y_x=max_ – Y_x=minimum_)/(*K*DApp + X), where × is the IgG or hACE2 concentration and Y is the normalized binding signal. Concentrations displaying hook effects, defined as concentrations higher than those generating the maximum PE MFI signal, were excluded from analysis. To maximize the dynamic range of potential differences in binding affinity to SARS-CoV-2 variants, binding experiments were conducted at each antibody’s respective SARS-CoV-2 *K*DApp concentration. Binding signal was normalized using the following equation: (MFI_anti-hu IgG PE_/MFI_anti-HA APC_) – (MFI_background anti-hu IgG_ PE/MFI_background anti-HA APC_), and calculated as a percentage of normalized signal of the reference WT SARS-CoV-2 strain RBD-SD1.

### ePCR library construction and selection of RBD mutants

SARS-COV-2 RBD-SD1 gBlock (IDT) was amplified by polymerase chain reaction (PCR) with iProof High-Fidelity PCR system (Bio-Rad, Cat # 1725310) following the manufacturer’s recommendations. The amplified DNA was purified (Nucleospin Gel and PCR Clean-up Kit, Macherey-Nagel, Cat # 740609.250) and subsequently mutagenized by error-prone PCR (ePCR) via the GeneMorph II Random Mutagenesis Kit (Agilent Technologies, Cat # 200550) with a target nucleotide mutation frequency of 0–4.5 mutations per kilobase of DNA. The mutagenized DNA product was cloned into yeast via electroporation as described earlier. The ePCR library was validated by plating a subset of the transformed ePCR yeast library on tryptophan dropout agar plates (Teknova, Cat # C6099) and sequencing single colonies. Prior to performing FACS selection, the ePCR RBD-SD1 library and WT RBD-SD1 yeast were induced as described above.

To select for mutants with diminished binding to ADG-2, induced cells were incubated for 30 minutes on ice with ADG-2 IgG diluted in PBSF to its EC_80_ concentration, which was calculated by titration on the yeast surface-displayed WT RBD-SD1 construct. Cells were washed twice in PBSF, stained in a secondary staining mixture, and analyzed on a BD FACS Aria II (Becton Dickerson), as described above. A subset of yeast population exhibiting HA-tag expression and reduced ADG-2 binding relative to the WT RBD-SD1 construct were sorted and propagated in SDCAA media for 48 hours at 30 °C. Selection was repeated for a second round to further enrich yeast encoding ADG-2 binding knock-down mutations. In the final round of selection, the induced library was stained with a mixture of recombinant hACE2-Fc, and S309 and CR3022 IgGs at their respective EC_80_ concentrations. The subset of the stained population that mirrored the binding profile of WT RBD-SD1-stained yeast was sorted and plated on agar plates for isolation and sequencing of single colonies. Clones possessing single amino acid substitutions identified from sequencing were cultured, induced, and evaluated for binding to ADG-2, S309, CR3022 IgGs and recombinant hACE2-Fc at their respective EC_80_ concentrations through flow cytometric analysis on the BD FACS Canto II (BD Biosciences). Binding signal was normalized and calculated as a percentage of the binding signal to reference WT RBD-SD1, as described above.

### Cryo-EM studies

SEC-purified SARS-CoV-2 HexaPro S was diluted to a concentration of 0.35mg/mL in a buffer composed of 2 mM Tris pH 8.0, 200 mM NaCl and 0.02% NaN_3_. Diluted spike was mixed with a two-fold molar excess of ADG-2 Fab and allowed to bind on ice for 5 minutes before the mixture was applied to a plasma-cleaned CF-1.2/1.3 grid. Excess liquid was blotted away using a Vitrobot Mark IV (Thermo Fisher) and the grid was vitrified by rapid plunging into liquid ethane. 4,748 micrographs were collected using Leginon (*45*) in a Titan Krios (Thermo Fisher) equipped with a K3 direct electron detector (Gatan). Motion correction, CTF-estimation and particle picking were performed in Warp (*46*) and extracted particles were imported into cryoSPARC v2.15.0 (*47*). 2D and 3D classification resulted in a final stack of 57,078 particles, which was used to calculate a 5.94 Å 3D reconstruction using non-uniform refinement (*48*). High-resolution crystallographic models of the SARS-CoV-2 RBD (PDB ID: 6M0J) (*31*) and a homologous Fab (PDB ID: 6APC) (*49*) were docked into the density using Chimera (*50*). A full description of the data collection and processing parameters can be found in Table S1.

### Animal studies

Twelve-month old female Balb/c mice (Envigo, strain 047) were treated with 200 μg of ADG-2 IgG via intraperitoneal (IP) injection at either 12 hours prior to infection (prophylactic) or 12 hours post-infection (therapeutic). Mice were anesthetized with ketamine/xylazine before being challenged with 1000 PFU of either SARS-CoV-MA15 or SARS2-CoV-2-MA10 (*34, 35*) via intranasal inoculation. Mouse body weights and respiratory function were monitored daily for 4 days. Respiratory function was monitored by whole body plethysmography (DSI) with a 30-minute acclimation period and a 5-minute measurement window as previously described (*51*). Viral lung titer was measured by plaque assay, assessing the lower lobe of the right lung. Gross pathology was performed on mice sacrificed on day 2 and day 4 post-infection. Gross pathology in the lung scored using a 4-point system, in which 0 represents no hemorrhage and 4 represents complete and total hemorrhage. All animal husbandry and experiments were performed at BSL3 and in accordance with all University of North Carolina at Chapel Hill Institutional Animal Care and Use Committee guidelines (AAALAC Institutional Number 329).

